# Structural basis of the dynamic human CEACAM1 monomer-dimer equilibrium

**DOI:** 10.1101/2020.07.14.199711

**Authors:** Amit K. Gandhi, Zhen-Yu J. Sun, Walter M. Kim, Yu-Hwa Huang, Yasuyuki Kondo, Daniel A. Bonsor, Eric J. Sundberg, Gerhard Wagner, Vijay K. Kuchroo, Gregory A. Petsko, Richard S. Blumberg

**Affiliations:** Division of Gastroenterology, Department of Medicine, Brigham and Women’s Hospital, Harvard Medical School, 75 Francis Street, Boston, MA 02115, USA.; Department of Cancer Biology, Dana-Farber Cancer Institute, Boston, MA 02215, USA.; Institute of Human Virology, University of Maryland School of Medicine, University of Maryland, 725 W Lombard St, Baltimore, MD 21201, USA.; Department of Biological Chemistry and Molecular Pharmacology, Harvard Medical School, 240 Longwood Avenue, Boston, MA, 02115, USA; Evergrande Center for Immunologic Diseases and Ann Romney Center for Neurologic Diseases, Harvard Medical School and Brigham and Women’s Hospital, 77 Avenue Louis Pasteur, Boston, MA 02115, USA.; Ann Romney Center for Neurologic Diseases, Department of Neurology, Brigham and Women’s Hospital, Harvard Medical School, Boston, MA 02115, USA..; Division of Gastroenterology, Department of Internal Medicine, Graduate School of Medicine, Kobe University, Kobe, 650-0017, Japan.; Department of Medicine, University of Maryland School of Medicine, University of Maryland, Baltimore, MD 21201,USA.; Department of Microbiology and Immunology, University of Maryland School of Medicine, University of Maryland, Baltimore, MD 21201, USA.; Department of Biochemistry, Emory University School of Medicine, Atlanta, GA 30322.

## Abstract

Human (h) carcinoembryonic antigen-related cell adhesion molecule 1 (CEACAM1) function depends upon IgV-mediated homodimerization or heterodimerization with host ligands, including hCEACAM5 and hTIM-3, and a variety of microbial pathogens. However, there is little structural information available on how hCEACAM1 transitions between monomeric and dimeric states which in the latter case is critical for initiating hCEACAM1 activities. We therefore mutated residues within the hCEACAM1 IgV GFCC’ face including V39, I91, N97 and E99 and examined hCEACAM1 IgV monomer-homodimer exchange using differential scanning fluorimetry, multi-angle light scattering, X-ray crystallography and/or nuclear magnetic resonance. From these studies, we describe hCEACAM1 homodimeric, monomeric and transition states at atomic resolution and its conformational behavior in solution through NMR assignment of the wildtype (WT) hCEACAM1 IgV dimer and N97A monomer. These studies reveal the flexibility of the GFCC’ face and its important role in governing the formation of hCEACAM1 dimers and potentially heterodimers.

## INTRODUCTION

Carcinoembryonic antigen-related cell adhesion molecule 1 (CEACAM1), also referred to as cluster of differentiation 66a (CD66a) and biliary glycoprotein (BGP), is a member of the carcinoembryonic antigen cell adhesion molecule (CEACAM) family of glycosylated immunoglobulin (Ig) molecules (*Kammerer & Zimmermann, 2010*). Expressed on the surface of several cell types, CEACAM1 has been demonstrated to play critical roles in morphogenesis (*Huang et al.,* 1999), apoptosis (*Nittka et al., 2004*), angiogenesis (*Ergun et al., 2000)*, cell proliferation (*Singer et al., 2010*) cell motility (*Ebrahimnejad et al., 2004*), fibrosis (*Satoh et al., 2017*), and most recently as an immunoreceptor important in mediating immune T cell tolerance (*Huang et al., 2015*). Human CEACAM1 (hCEACAM1) is a single pass type I transmembrane protein expressed as 12 alternatively spliced isoforms that all contain an N-terminal V set fold of the immunoglobulin superfamily (IgV) ectodomain followed by up to three type 2 constant immunoglobulin (IgC2) ectodomains (A1, B, A2), a transmembrane sequence, and a signaling cytoplasmic domain. Depending on splice variation, the cytoplasmic domain either includes a long (L) sequence inclusive of two immunoreceptor tyrosine-based inhibitory motifs (ITIMs) or a short (S) domain devoid of ITIMs (*Beauchemin et al., 1999*) that impart intracellular inhibitory or non-inhibitory signals, respectively.

CEACAM1 function is triggered by intercellular or *trans* binding of the N-terminal IgV domain, resulting in higher order surface CEACAM1 oligomerization and subsequent intracellular signal transduction. In contrast to other immunoreceptors such as the T cell inhibitory and mucin domain containing protein 3 (TIM-3), programmed cell death protein 1 (PD-1), programmed death-ligand 1 (PD-L1), and cytotoxic T-lymphocyte-associated protein 4 (CTLA-4), CEACAM1 serves as its own primary ligand owing to high affinity homophilic interactions of its unique IgV domain and is an important microbial receptor (*Kim el al., 2019*). At basal steady state, CEACAM1 alternates between monomeric and *cis* homodimeric forms on the cell surface (*Klaile et al., 2009*), thus presenting a conundrum for *trans* interactions due to the requirement of an accessible CEACAM1 monomer and more specifically, an exposed IgV domain for ligand binding. Therefore, CEACAM1 must undergo a dynamic process of *cis* monomer-dimer exchange and *trans* dimer-higher order oligomerization for productive CEACAM1 activation. At present, the structural details of the monomer-dimer-higher order oligomer exchange mechanism are not well understood.

The hCEACAM1 IgV domain contains 108 amino acids arranged in 9 beta strands (ABCC’C”DEFG) that fold into the conserved IgV anti-parallel beta-sandwich tertiary structure adopted by other IgV-containing immunoreceptors including TIM-3 (*Fedarovich, Tomberg, Nicholas, & Davies, 2006; Huang et al., 2015; Huang et al., 2016)*, PD-1 and PD-L1 (*Lin et al., 2008*). The opposing ABED and GFCC’ faces of the CEACAM1 beta-sandwich are tethered by an internal salt bridge (R64:D82) that mimics a stabilizing covalent disulfide linkage found in most Ig domains (*Huang et al., 2015)*. Although the ABED surface is exclusively glycosylated, CEACAM1 has been suggested to exist in diverse oligomeric states (*Bonsor, et al., 2015)* that include an ABED-mediated homodimer (*Fedarovich et al., 2006*), but the more dominant oligomeric form appears to be the high-affinity GFCC’-mediated homodimer (*Bonsor et al., 2015; Huang et al., 2015*) that shares a conserved quaternary structure to other IgV domain-containing proteins. Unique to the hCEACAM1 IgV GFCC’ surface is the prominent protrusion of the CC’ loop that differs significantly from the ordered β-hairpin observed in other described IgV structures. The displaced CC’ loop forms a cleft with the FG loop that exposes the key residues F29, S32, Y34, V39, G41, Q44, Q89, I91, N97, and E99 critical in mediating homophilic CEACAM1 interactions *(Bonsor et al., 2018; Fedarovich et al., 2006; Huang et al., 2016; Huang et al., 2015*). As demonstrated in our previously reported crystal structure (PDB code 4QXW) of the CEACAM1 homodimer (*Huang et al., 2015*), the side chains of residues S32, Y34, Q44, Q89, N97, and E99 form the hydrogen bonding network at the GFCC’ interface and additional side-chain to main-chain backbone interactions between S32 to L95, Q44 to N97, and E99 to G41 and hydrophobic interactions by residues F29, V39, and I91 further coordinate the GFCC’-mediated homodimer (***Figure 1***-***Figure 1 supplements A-B***). However, while these residue-level interactions determine specificity of the homodimerization interface, they have been recently reported to impart varied free energy contributions to the strength of the interaction (*Bonsor et al., 2018*), raising intrigue in the underlying mechanisms of CEACAM1 IgV monomer-dimer exchange.

**Figure 1.**
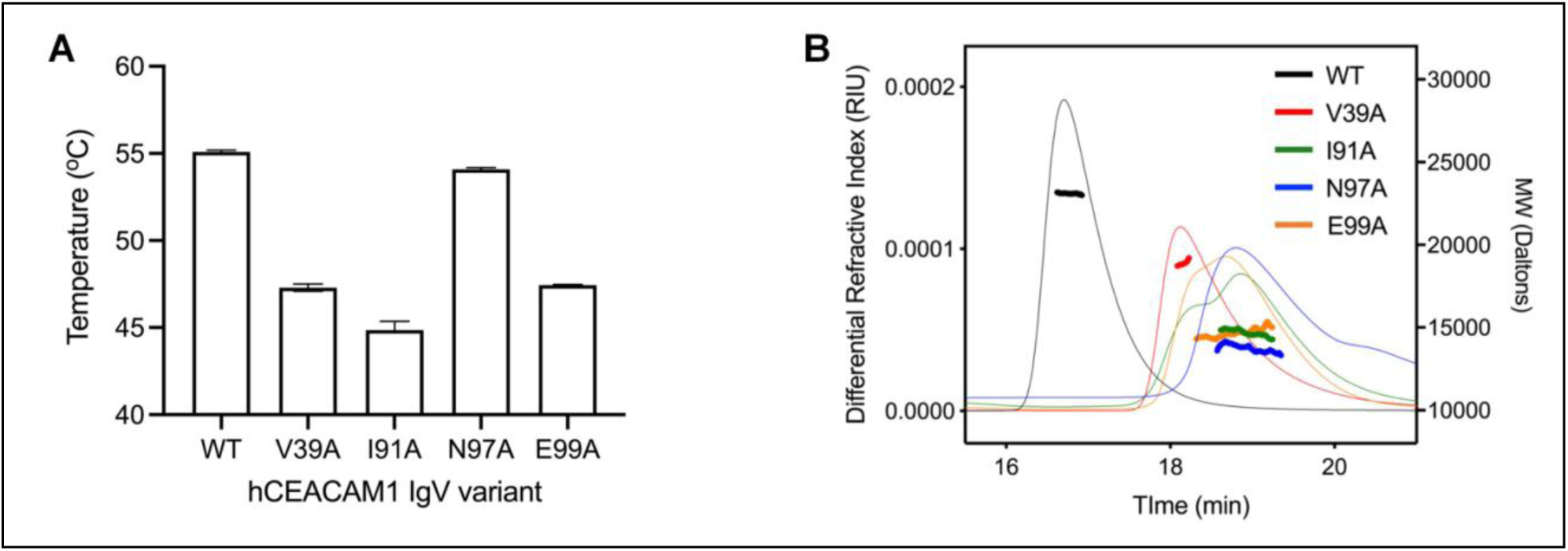
Biophysical characterization of CEACAM1 IgV mutants. Thermal stability and molecular size analysis of hCEACAM1 WT and GFCC’ face mutants. (A) Variations in melting point temperature (*T_M_*) determined by differential scanning fluorimetry (DSF) are shown for WT and mutant hCEACAM IgV. (B) Size exclusion chromatography and multi-angle light scattering (SEC-MALS) differential refractive index (RIU) chromatograms and calculated molecular weights are displayed for WT (black), V39A (orange), I91A (green), N97A (blue) and E99A (red). **Figure supplement 1.** Crystal structure of the IgV of hCEACAM1 homodimer (PDB code 4QXW).

Although the major mode of CEACAM1 binding is homophilic, several other host and microbial ligands also exist. The binding of cell surface CEACAM1 by these ligands induces higher order multimerization and in the case of microbial ligands, hijacks the downstream signaling machinery to achieve survival gain. Surprisingly, all of the described CEACAM1 host and microbial ligand interactions involve the GFCC’ surface on CEACAM1, thereby requiring disruption of CEACAM1 homophilic interactions to allow for participation in the heterophilic interactions. The IgV domain of hCEACAM1 has high sequence similarity with hCEACAM family members (***Supplements table 1A-C***), although only CEACAM5, commonly referred to as CEA, appreciably binds to CEACAM1 owing to conservation of residues on its respective GFCC’ surface, including the CEACAM1-homodimerization dependent residues F29, S32, V39, R43, Q44, I91 and E99 (*Bonsor et al., 2015; N Korotkova et al., 2008)*. More recently, biochemical (*Y. Huang et al., 2015)* and NMR studies (*Gandhi et al., 2018; Y.-H. Huang et al., 2016*) demonstrated calcium-dependent binding of the N-terminal IgV domain of TIM-3 to CEACAM1 also through GFCC’-mediated interactions.

Despite the requirement of competing with CEACAM1 as a ligand, several bacteria, fungi, and a virus that include *E. coli* (*Korotkova et al., 2008*), *Neisseria sp.* (*Watt et al., 1997*), *Moraxella catarrhalis* (*Conners et al., 2008)*, *Haemophilus influenza (Edwards et al., 2005*), *Helicobacter pylori* (*Königer et al., 2016*), *Fusobacterium sp.* (*Brewer et al., 2019*), *Candida sp.* (*Klaile et al., 2017)*, and the coronavirus murine hepatitis virus (*Tan et al., 2002)*, have evolved structurally distinct microbial receptors that universally disrupt CEACAM1 homophilic interactions at the GFCC’ surface to form unique heterodimeric interactions (*Tchoupa et al., 2014)*. Recently, biochemical studies on the *H. pylori* surface protein HopQ demonstrated that HopQ effectively disrupts the hCEACAM1 IgV homodimer by outcompeting homophilic hCEACAM1 IgV GFCC’ surface interactions (K_D_ = 450 nM) through up to 20-fold higher affinity heterophilic interactions (K_D_ = 23-279 nM) targeting the GFCC’ surface (*Bonsor et al., 2018; Moonens et al., 2018*). The crystal structure of the hCEACAM1 IgV-HopQ complex demonstrated direct involvement of the GFCC’ surface on hCEACAM1 in complex formation; however, there were significant differences in the structural and biophysical features that distinguished hCEACAM1 IgV-HopQ heterodimerization from hCEACAM1 homodimerization (*Bonsor et al., 2018*). One specific discriminating feature involves residue N97, which has been reported to nearly abrogate hCEACAM1 homodimerization (K_D_ ∼ 1 mM) but does not significantly affect HopQ binding (*Bonsor et al., 2018*). This observation raises the question about the role of specific residues at the GFCC’ face in determining hCEACAM1 homophilic and heterophilic interactions, but just as importantly emphasizes the need to decipher the underlying structural and biochemical features that determine hCEACAM1 monomer-homodimer exchange, which enables the formation of these various interactions. At present, the role of the GFCC’ face and specific residues in determining the basal monomer-dimer equilibrium at steady-state and moreover the conformational behavior of hCEACAM1 in solution still remain elusive.

Here we describe the structural and biochemical features of specific hCEACAM1 mutants present at the GFCC’ surface (V39A, I91A, N97A, E99A) in static conformation by x-ray crystallography and in solution by nuclear magnetic resonance (NMR). The unique crystal structures reveal a range of subtle and gross conformational changes in the CEACAM1 homodimer interface that highlight the significance of each examined residue. Furthermore, dynamic NMR studies of the N97A mutant substantiates the critical role of this residue in mediating CEACAM1 homodimerization. These studies illuminate the mechanisms that govern dynamic CEACAM1 homodimerization and provide a foundation for exploring how foreign ligands exploit CEACAM1 as well as serving as a basis for therapeutic intervention.

## Results

### 1. Biophysical characterization of the hCEACAM1 IgV GFCC’ face mutants

In order to probe the role of the GFCC’ surface in determining the monomer-dimer equilibrium, we introduced alanine substitutions at residues V39, I91, N97, and E99, which have been described to be important for CEACAM1 IgV homodimerization and identified as naturally occurring single nucleotide polymorphisms (SNPs) sites [rs772794650 (I91M), rs1335884800 (N97T), and rs142826356 (E99G)]. We first expressed and purified WT and site-specific mutant hCEACAM1 IgV proteins using our published protocols (*Huang et al., 2016*) and measured variations in their respective thermal denaturation temperature (*T_M_*), reflective of their stability by differential scanning fluorimetry (DSF) (***Figure 1A***). We observed a single *T_M_* for each protein at 25 μM, suggestive of a single step denaturation event despite whether the protein was expected to be a hCEACAM1 IgV monomer (N97A) or dimer (WT). There was also a correlation of decreasing *T_M_* with decreasing homodimerization affinity of the different hCEACAM1 mutants, suggesting that weakening of hCEACAM1 IgV homodimerization destabilizes hCEACAM1 IgV stability. However, the hCEACAM1 IgV N97A variant that has been reported to be monomeric, with a K_D_ approaching 1 mM (*Bonsor et al., 2018*), exhibited a similar melting temperature (54.09 °C) compared to WT protein (55.09 °C), suggesting a unique stabilizing property of an alanine at that position and/or promotion of a monomeric state. Next, we assayed the solution characteristics of each hCEACAM1 IgV sequence variant by analytical size exclusion chromatography and multi-angle light scattering (SEC-MALS) and calculation of absolute molecular weight. Each hCEACAM1 IgV variant (100 μM) eluted as a single peak but with varying calculated molecular weight ranging from dimer (WT, 23.1 kDa) to monomer (N97A, 13.5 kDa) (***Figure 1B***). The presence of a single peak for each protein variant and varying intermediate absolute molecular weights suggests rapid rates of exchange between monomeric and dimeric states of the IgV domain rather than a slow equilibrium compared with the time scale of the experiments.

### 2. High resolution crystal structures of hCEACAM1 IgV GFCC’ face mutants

To further determine the impact of V39, I91, N97, and E99 on hCEACAM1 homodimer formation and probe their contributions to the hydrophobic and hydrogen bonded network at the GFCC’ face, we solved the crystal structures of individual V39A, I91A, N97A, and E99A mutant proteins to 1.9, 3.1, 1.8, and 1.9 Å resolution, respectively. The I91A and E99A mutants were crystallized in high tascimate buffer conditions with the symmetry of a tetragonal space group, which were very similar to hCEACAM WT IgV (PDB code 4QXW) crystallization conditions and resolved space group, however with varied unit cell constants (***Table 1***). The V39A and N97A mutants were crystallized in conditions different from high tascimate buffer and had the symmetry of unique trigonal and c-centered orthorhombic space groups, respectively (***Table 1***).

The crystal structures of the I91A and E99A mutants revealed a globally similar GFCC’ face dimer (***Figures 2A-2D***), as observed for the hCEACAM WT dimer (PDB code 4QXW) but with various localized conformational differences resulting in a C-alpha root mean square deviation (RMSD) of 0.657 Å (over 1489 atoms) and 0.305 Å (over 1539 atoms) for the I91A and E99A mutants, respectively (***Figure 2, Figure 2-figure supplement 4A***, ***Figure 2-figure supplement 5A***). More interestingly, for both of the I91A and E99A mutants, although a homodimer was observed in the crystal asymmetric unit, weaker hydrogen bonded and hydrophobic interactions, necessary for homodimer formation through the GFCCC” face, were observed between two hCEACAM1 molecules compared to WT hCEACAM1 (***Figures 2A-2D***, ***Figure 1***-***Figure 1 supplements A-B***, ***Figure 2-figure supplements 2A-2D***, ***Figure 2-figure supplements 3A-3D***). For the I91A mutant (***Figures 2A-B***), specifically, the hydrogen bonded interactions between the residues N97-*Y34* (using nomenclature convention here and after, where N97 residue is from molecule (a) and *Y34* residue in italics is from (*b*) molecule present in the crystal asymmetric unit), Q89-*Y34,* S32-*N97, Y34-N97,* and Q44-*N97* were disrupted (***Figure 2B***, ***Figure 2-figure supplement 2B***).

**Figure 2.**
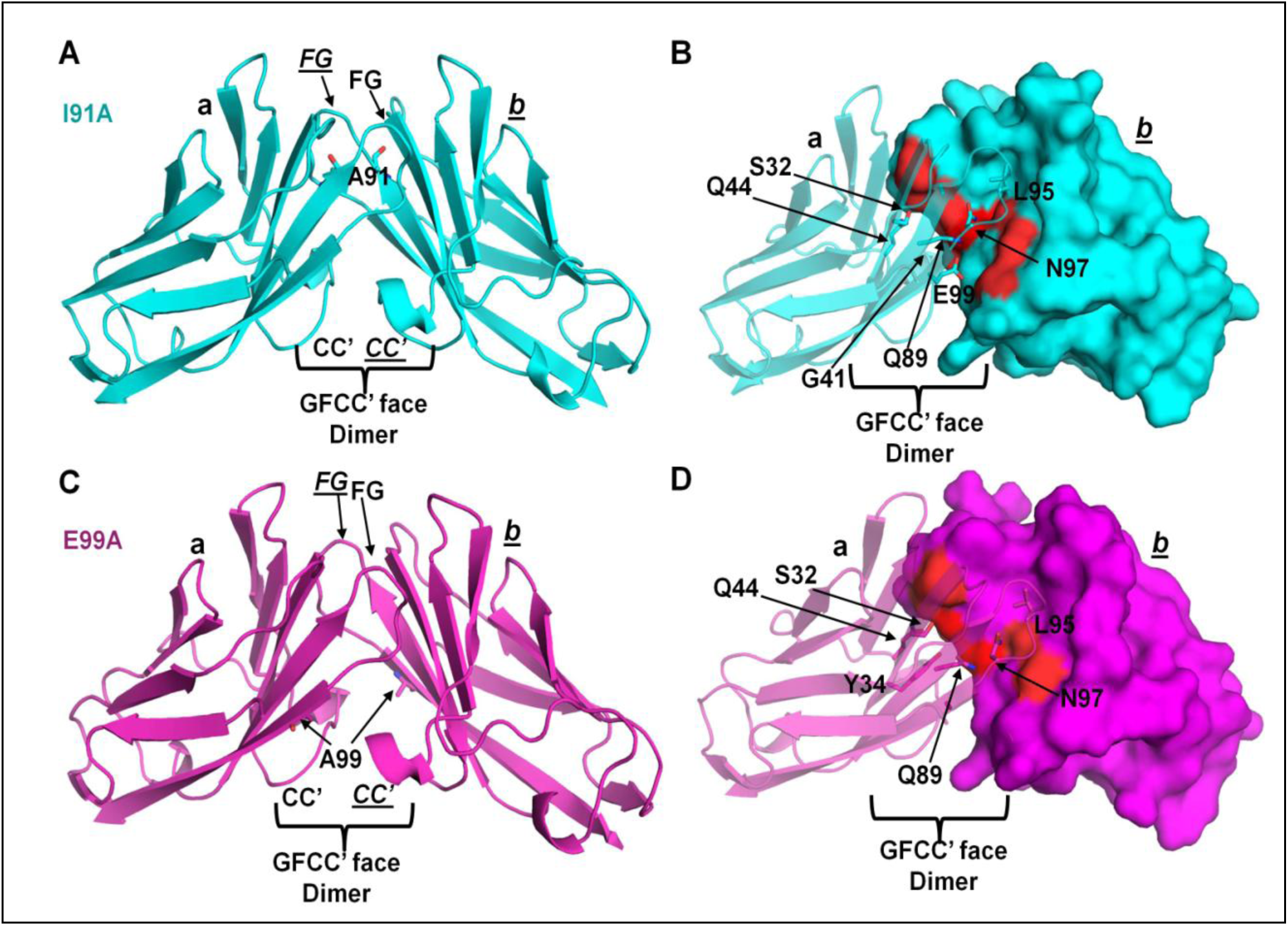
Crystal structures of the I91A and E99A IgV mutants of hCEACAM1. (A) Ribbon diagram (cyan) of the molecule (a) and molecule (*b*) as observed in the unit cell of the I91A crystal structure. The A91 residues of molecule (a) and molecule (*b*) are shown by stick representation. The FG and CC’ loops that mediates formation of GFCC’ face dimer are labeled in bold for the molecule (a) and labeled in italics and underlined for the molecule (*b*). (B) The ribbon and surface diagram of the molecule (a) and (*b*) in cyan as observed in the I91A crystal structure. The S32, G41, Q44, Q89, L95, N97, and E99 residues that mediates hydrogen bonded interactions in the formation of GFCC’ face dimer of I91 crystal structure are shown by stick and surface bright and light red representations for molecule (a) and (*b*), respectively. Specifically, the molecule (*b*) residues that make multiple hydrogen bonded interactions across the GFCC’ face are shown by bright red surface representation. (C) Ribbon diagram (magenta) of the molecule (a) and molecule (*b*) as observed in the unit cell of the E99A crystal structure. The A99 residues of molecule (a) and molecule (*b*) are shown by stick representation and FG and CC’ loops are labeled as described above. (D) The ribbon and surface diagram of the molecule (a) and (*b*) in magenta as observed in the E91A crystal structure. The S32, Y34, Q44, Q89, L95, N97, and E99 residues that mediates hydrogen bonded interactions in the formation of GFCC’ face dimer of I91 crystal structure are shown by stick and surface bright and light red representations as described above. **Figure supplement 2**. Structural comparison of the I91A mutant and hCEACAM1 WT IgV (PDB code 4QXW) crystal structures. **Figure supplement 3**. The E99A mutant crystal structure compared with the hCEACAM1 WT IgV (PDB code 4QXW) crystal structure.

Further, the hydrophobic interactions between two F29 residues were weaker due to conformation changes and increased distance compared to WT (***Figure 2B***, ***Figure 2-figure supplements 2C-D***). Thus, the weakening of the hydrogen bonded and hydrophobic interactions highlighted the importance of hydrophobic interactions that two I91 residues of hCEACAM1 molecules provide in the formation of other hydrogen and hydrophobic interactions at the GFCC’ face. Similarly, for the E99A mutant (***Figures 2C-D***), mutation of glutamic acid at position 99 to alanine abrogated side chain to main-chain backbone interactions between E99-*G41* of both hCEACAM1 molecules as expected, but also abrogated the hydrogen bonded interactions between Q89-*Y34, N97-Y34, and Q89-N97* residues of two hCEACAM1 molecules (***Figure 2D***, ***Figure 2-figure supplement 3B***). In addition, the distance between two hydrophobic V39 residues was slightly higher as measured by a distance of 3.9 Å between V39-β carbons compared to distance of 3.7 Å in the WT (***Figure 2D***, ***Figure 2-figure supplements 3C-D***). Thus, the loss of important hydrogen bond interactions and possibly weaker hydrophobic interactions observed in the I91A and E99A crystal structure support the weak dimeric nature of these mutants as observed in our biophysical studies and previously reported mutagenesis studies (*N Korotkova et al., 2008; Watt et al., 2001)*.

The diffraction data sets of various V39A mutant crystals showed twinning and one of the resolved crystal structures from the single high resolution data set (1.9 Å) revealed two copies of an hCEACAM1 GFCC’ face dimer (***Figure 3A***) with RMSD of 0.677 Å (over 1482 atoms) and 3.077 Å (over 1655 atoms) for the first V39A dimer (***Figure 3-figure supplement 5A***, formed by molecules (a) and (*b*) and second V39A dimer (***Figure 3-figure supplement 4A***, formed by molecules c and (*d*)), respectively. The first dimer resembled a WT GFCC’ face dimer with minor conformational changes (***Figure 3-figure supplement 5A***). More interestingly, the second GFCC’ face dimer comprising molecules (c) and (*d*) showed significant conformational differences across various strands and loops (***Figure 3-figure supplement 4A***), as shown by increased distances between the interacting CC’ loops (10 Å between β carbon of V39 residue compared to distance of 3.7 Å between β carbon of V39 in the WT) (***Figure 3-figure supplement 4A***). Further, we observed significantly weaker and a decreased network of hydrogen bonded interactions within the FG and CC’ loop residues in the weak V39A dimer (***Figure 3B, Figure 3-figure supplement 4B***) compared to hCEACAM1 WT GFCC’ face dimer (PDB code 4QXW) (***Figure 3-figure supplements 6A-6B***). Specifically, we observed abrogation of all hydrogen bonded interactions mediated by residues S32, G41, L95, and E99 of (c) molecule and residues S32, Y34, G41, Q89, and E99 of (*d*) molecule. (***Figure 3B, Figure 3-figure supplement 4B***). In addition, N97 residues of c and (*d*) molecules mediated only two hydrogen bonded interactions compared to seven hydrogen bonded interactions observed in hCEACAM1 WT homodimer (***Figure 3-figure supplement 4B, Figure 3-figure supplement 6A***). Further, we observed weaker hydrophobic interactions compared to WT (***Figure 3-figure supplements 4C-D, Figure 3-figure supplement 6B***) as revealed by conformational changes and an increased interaction distance of residues F29 and I91 residues of molecules (c) and (*d*). Residue V39 and residues F29 and I91 mediate hydrophobic interactions necessary for the stabilization of the GFCC’ face dimer as observed in the hCEACAM1 homodimer (PDB code 4QXW) (***Figure 3-figure supplement 6B)***. Thus, the weak V39A GFCC’ face dimer demonstrated significantly weaker CC’ and FG loop interactions, consistent with a less stable and interacting GFCCC” face that possibly reflects one of the transition states associated with exchange between a hCEACAM1 monomer and dimer. Similar weak V39A GFCC’ face dimer was observed in the resolved crystals from the two additional data sets (Not reported). Interestingly for both of the V39A dimers, a significant metal ion density at the 4.0 σ level in a 2Fo-Fc electron density map was observed, which we refined for Nickel (Ni^++^). Here, we observed hexa-coordination of Ni^++^ by residues H105 and V39 from three neighboring hCEACAM1 molecules in the unit cell and crystallographic symmetry mates (***Figure 3-figure supplements 5B-C***). Overall, the V39A crystal structure demonstrated a significantly weaker GFCC’ face dimer and provided the structural basis for the monomer-inducing feature of the V39A substitution suggested previously *(N Korotkova et al., 2008; Watt et al., 2001*).

**Figure 3.**
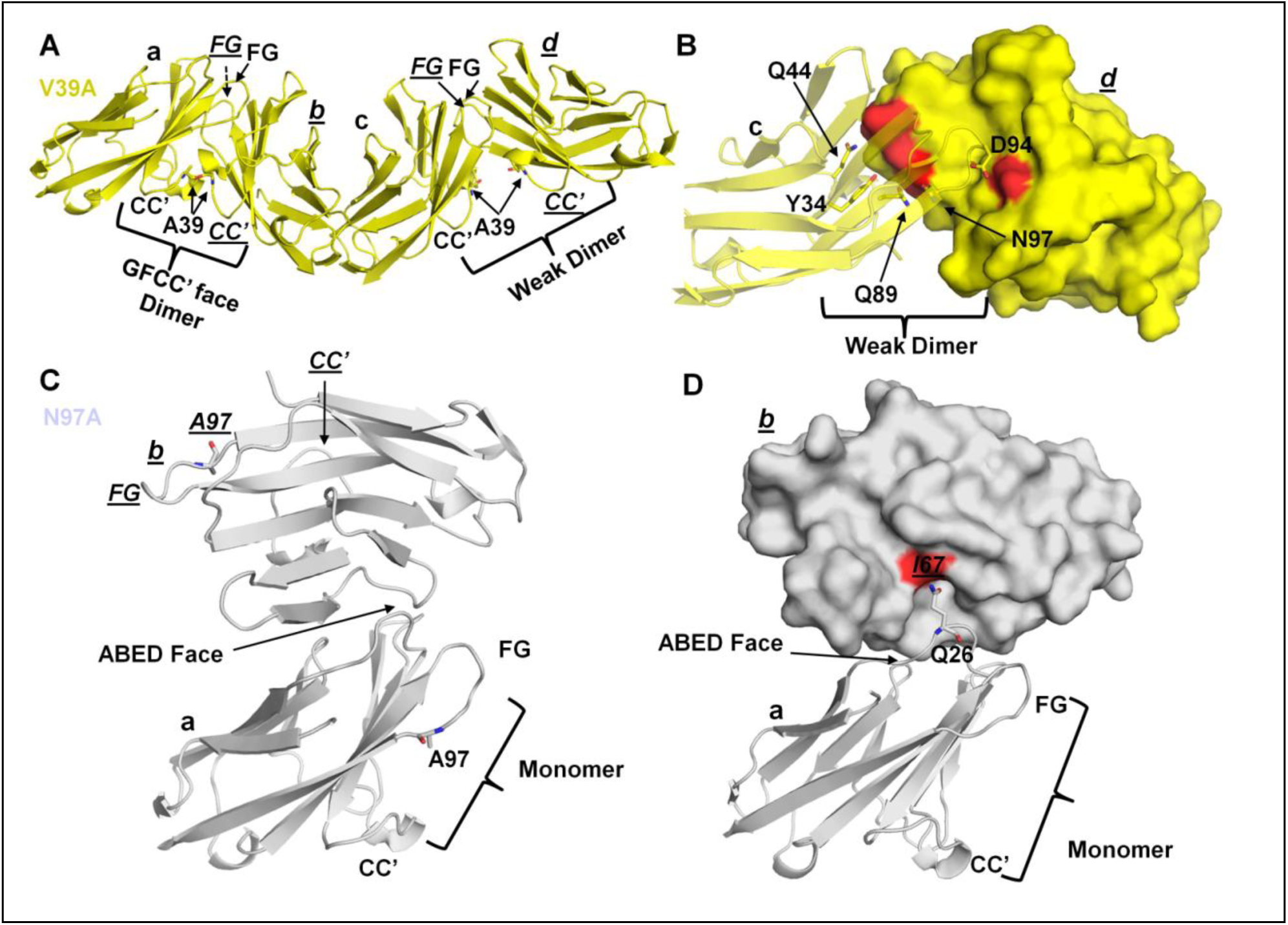
Crystal structures of the V39A and N97A IgV mutants of hCEACAM1. (A) Ribbon diagram (yellow) of the molecules (a), (*b*), (c), and (*d*) as observed in the unit cell of the I91A crystal structure. The V39 residues of these four molecules are shown by stick representation and the FG and CC’ loops are labeled in bold for the molecules (a, c) and labeled in italics and underlined for the molecule (*b, d*). The molecules (a) and (*b*) make dimer that mimic GFCC’ face dimer as observed in the hCEACAM1 (PDB code 4QXW) crystal structure (Figure *1 supplement).* The molecules (c) and (*d*) make weak GFCC’ face dimer where FG and specifically CC’ loops are far apart and very few GFCC’ face residues mediates the interactions. (B) The residues Y34, Q44, Q89, D94 and N97 that mediates formation of weak GFCC’ face dimer are shown in stick and surface bright and light red representations for molecules (c) and (*d*) in yellow, respectively. (C) The crystal structure of N97A mutant with two monomeric molecules (a, bottom) and (*b*, top) shown by ribbon diagram colored silver white. The A97 residues of both molecules are shown by stick representation and this mutation leads to abrogation of GFCC’ face dimer in the N97A crystal structure. The CC’ and FG loops are labeled and very limited interface contact between two N97A molecules through ABED face is shown by arrow. (D) The ribbon and surface diagram of N97A monomeric molecules (a) and (*b*), respectively, as observed in the unit cell of N97A crystal structure. The residues Q26 (stick representation) of molecule (a) and I67 (surface colored red) of molecule (*b*) make two hydrogen bonded interactions and form minor point of contact between two molecules at ABED face. **Figure supplement 4**. The structural basis of weak V39A mutant GFCC’ face dimer formation and comparison with hCEACAM1 WT IgV (PDB code 4QXW) crystal structure. **Figure supplement 5**. V39A mutant dimer formed by molecules (a) and (*b*) mimic hCEACAM1 WT GFCC’ face dimer (PDB code 4QXW) and residues H105 and V106 with their symmetry mates mediates binding with Nickel (Ni **^++)^**. **Figure supplement 6**. Hydrogen bonds and hydrophobic interactions of the IgV of hCEACAM1 homodimer (PDB code 4QXW) and N97-mediated asymmetrical hydrogen bonded interactions.

Another feature of the homodimeric interface was the strong propensity of N97 to mediate a hydrogen bond network comprised of seven hydrogen bond interactions in the WT homodimer crystal structure (PDB code 4QXW) (***Figure 3-figure supplements 6A***). It is important to note that the interactions mediated by N97 on molecules (a) and (*b*) are not perfectly symmetrical. Specifically, whereas the N97 residue of molecule (a) was observed to interact with S32, Y34, and Q44 residues of molecule (*b*), N97 of molecule (*b*) interacted with residues S32, Y34, Q44, and additionally residue Q89 of molecule (*a*) (***Figure 3-figure supplements 6C***). To probe the importance of N97-mediated asymmetrical hydrogen bonded interactions in the formation of a WT homodimer, we examined the N97A crystal structure, and observed two N97A molecules (a) and (*b*) in the crystal asymmetric unit with overall similar structure fold compared to a single hCEACAM1 molecule of the hCEACAM WT dimer (PDB code 4QXW) (***Figures 3C-D***). Importantly, the GFCC’ face dimer was completely absent in the N97A crystal structure even at the crystallographic concentration that exceeded 800 μM. This abrogation of the GFCC’-mediated dimer was consistent with previous studies (*Bonsor et al., 2018*) and the results of our SEC-MALS data, which showed the N97A mutant to be a monomeric protein in solution (***Figure 1B***). Although the N97A mutant crystallized with two hCEACAM1 molecules in the crystal asymmetric unit with an RMSD of 0.454 Å (over 638 atoms) and 0.561 Å (over 628 atoms) compared to a single hCEACAM1 molecule present in the homodimer structure (PDB code 4QXW), only two hydrogen bonded interactions below 3.5 Å distance were observed between the two molecules of N97A, involving residues G26 of molecule (a) and I67 of molecule (*b*) (***Figures 3C-D***). Thus, the hCEACAM1 IgV N97A crystal structure highlighted that substitution of asparagine at position 97 to alanine eliminated a crucial hydrogen bond network mediating hCEACAM1 IgV dimerization resulting in abrogation of higher order oligomerization (PDB code 4QXW) (***Figure 3-figure supplements 6A-C***). To further confirm a monomeric state of the N97A mutant as observed in the crystal structure, we carried out PDB-PISA (Protein Interfaces, Surfaces and Assemblies) analysis, which computes a complex significance score (CSS) of macromolecular complex formation by accounting for the energy of complex formation, interface area, number of bonds formed, and hydrophobicity variables with solvation energy gain (*Krissinel & Henrick, 2007*) (***Supplementary Tables 2, 3***). We observed a CSS score of 0.0 for the two monomers present in the N97A crystal structure compared to a CSS score of 0.89 for the hCEACAM1 WT GFCC’ face dimer (***Supplementary Tables 3***). We also determined the CSS scores of other N97A molecules present in the crystallographic symmetry unit and found that no significant CSS score was observed compared to the hCEACAM1 GFCC’ face dimer. Thus, the PDB-PISA analysis confirmed abrogation of a GFCC’ face dimer in the N97A crystal structure and provided a structural basis for the critical role of the N97 residue in determining the monomer and dimer state of hCEACAM1.

Further, we extended the PDB-PISA analysis to the V39A, E99A, and I91A mutants as well (***Supplementary Table 3***). Consistent with SEC-MALS data reflecting the monomerizing nature of V39A protein in solution and rapid monomer-dimer exchange, we observed a CSS score of 0.0 for the weak GFCCC” face V39A dimer (formed by molecules c and *d*) and CSS score of 1.0 for the V39A dimer formed by molecules (a) and (*b)* that resembled the hCEACAM1 WT homodimer (***Supplementary Table 3***). Similarly, we observed a CSS score of 0.631 for the E99A mutant, which demonstrated the presence of a slightly weakened GFCC’ face dimer and highlighted the role of an E99 side chain to G41 main-chain backbone interaction in the formation of a hCEACAM1 homodimer. Interestingly, PDB-PISA analysis indicated a CSS score of 1.0 for the I91A dimer, which possibly reflects an overall conservation of hydrogen bonded and hydrophobic interactions consistent with the observed homodimer in the I91A crystal structure. Overall, our high resolution crystal structure studies demonstrated atomic level contributions from V39, I91, N97, and E99 within the GFCC’ face to hCEACAM1 homodimer formation. Further, they reveal that whereas the V39A mutant crystallized as a WT-weakened dimer configuration consistent with a transition state through effects on the CC’ loop, the N97A mutant mimics a monomeric form of hCEACAM1.

### 3. NMR structural studies of the hCEACAM1 dimer and N97A mutant

To probe the role of the GFCC’ face and N97A mutation in determining the dimerization characteristics of hCEACAM1 in solution, we performed solution NMR spectroscopy studies using isotopically labeled WT and N97A mutant-containing hCEACAM1 IgV proteins (***Figures 4A-B, 5A-5B***, ***Figure 4-figure supplement 7A-B***). We first purified ^15^N/^13^C double labeled tagless hCEACAM1 IgV WT protein using our published protocols (*Huang et al., 2016*). During the time course of the triple resonance experiments, the NMR signal intensities decreased, suggesting the dynamic formation of high order oligomer and/or soluble aggregates without any observed precipitation. A ^15^N/^13^C/^2^H triply labeled hCEACAM1 protein sample was needed to acquire the full non-uniformly sampled (NUS) *(Robson et al., 2019)* data set that enabled us to complete 100% of the NMR backbone assignments for the hCEACAM1 WT IgV dimer (***Figure 4-figure supplement 7A***), compared to the previously published 70% assignments (*Zhuo et al., 2016*). The secondary structures of the hCEACAM1 WT IgV were predicted from the assigned NMR chemical shift values using the TALOS-N program (*Shen & Bax, 2013*), which agree well with secondary structures (*Huang et al., 2015*) observed in the dimeric crystal structure (PDB code 4QXW) of the hCEACAM1 IgV (***Figure 4-figure supplement 8A***). It should be noted that residue G41 has a remarkable downfield shifted ^15^NH peak position in the ^15^N-HSQC spectrum (***Figure 4-figure supplement 7A***), which is typically associated with strong hydrogen-bond formation. Indeed, in the crystal structure of hCEACAM1 WT dimer, the backbone amide of residue G41 forms an inter-molecular hydrogen bond with the side chain carboxy group of residue E99 (***Figure 1-figure supplement 1A, Figure 3-figure supplement 6A***). Thus, Gly41 provided us with an easily identifiable NMR indicator for hCEACAM1 IgV dimer formation.

**Figure 4.**
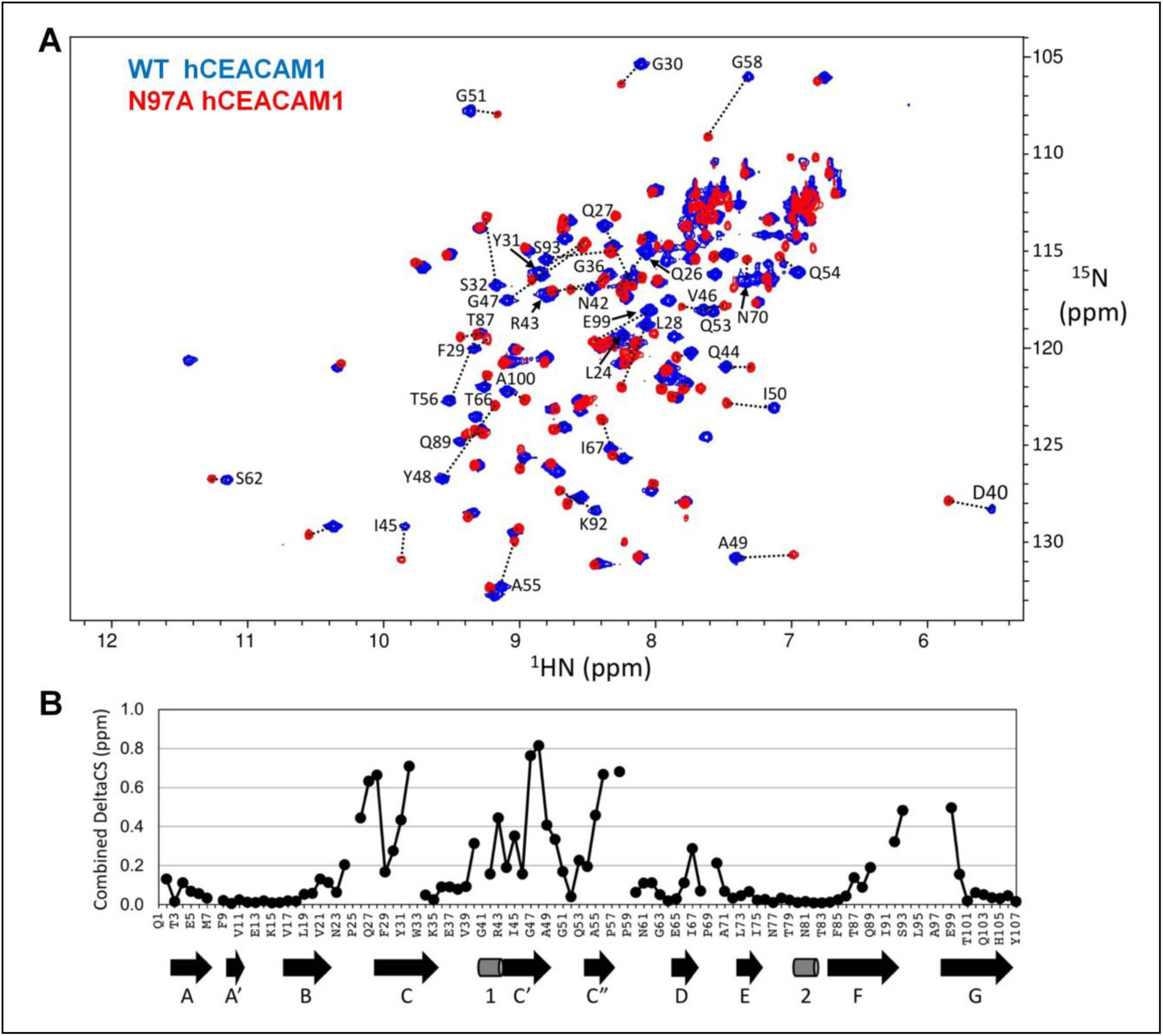
(A) Overlaid ^15^N-HSQC spectra of WT (blue) and N97A mutant (red) hCEACAM1 IgV protein. The corresponding assigned residues with significant peak shifts were indicated and connected by dotted lines. (B) Combined ^1^HN and ^15^N chemical shift changes, sqrt (csH^2^ + csN/5)^2^, between WT and N97A mutant hCEACAM1 IgV are shown in comparison with the secondary structure elements from the X-ray structure of hCEACAM1 WT depicted below. **Figure supplement 7** ^15^N HSQC spectra of hCEACAM1 IgV WT and N97A mutant proteins. **Figure supplement 8** Secondary structures prediction and comparison. **Figure supplement 9** Mapping of NMR chemical shift changes.

Next, we purified ^15^N/^13^C double-labeled N97A mutant-containing hCEACAM1 IgV protein to determine the behavior of the N97A-containing domain in solution. The ^15^N-HSQC spectrum of the N97A mutant showed large-scale spectral changes for nearly a third of the backbone amide groups compared to the WT IgV domain (***Figure 4A***), even though only a single asparagine residue was mutated to alanine. The molecular size of the N97A protein was estimated by TRACT (TROSY for RotAtional Correlation Times) measurements (*Lee, Hilty, Wider, & Wüthrich, 2006)*. The average molecular rotational correlation time was determined to be 6.5 ns on a 500MHz spectrometer using a freshly prepared 100 μM, ^15^-N N97A samples, corresponding to a ∼11 kD molecular weight monomer. Thus, a monomeric nature of N97A mutant was confirmed in solution and was consistent with abrogation of the GFCC’ face in the crystal structure, PDB PISA analysis and results of biophysical characterization studies described above.

However, the NMR signal intensities of roughly half the N97A mutant protein peaks continued to decay at room temperature, while the remaining peaks only showed minor shifts, suggesting that N97A mutant hCEACAM1 protein could also slowly form high order oligomer and/or soluble aggregates. The rotational correlation time as measured by TRACT experiment increased to 18 ns in an 8-day old 100 μM protein sample maintained at 25°C. These observations are consistent with the results of a previous study that examined the dynamic oligomer to monomer shift of hCEACAM1 by inside out signaling and hinted at the strong propensity of monomeric CEACAM1 to form oligomers (*Patel et al., 2013*). Even though we observed poor NMR signals of some of the resonance peaks, we were able to complete 90% of the backbone amide resonance assignments for the N97A mutant (***Figure 4-figure supplement 7B***). A ^15^N/^13^C/^2^H triple-labeled N97A mutant hCEACAM1 protein sample was produced to acquire additional NMR data sets for backbone assignments. However, per-deuteration of protein that reduced NMR relaxations and increased NMR signals for the WT hCEACAM1 made little improvement in the N97A mutant assignments (the reason for which will be discussed below).

The overall secondary structures of the N97A mutant as predicted by the assigned NMR chemical shifts were similar to those of the WT protein (***Figure 4-figure supplement 8B***), and also consistent with secondary structures observed in the crystal structure of the N97A mutant. The largest backbone amide NMR chemical shift changes for the N97A mutant relative to the hCEACAM1 WT were found among residues in or near the GFCC’ face, the homodimer interface for the WT protein (***Figure 4B***, ***Figure 4-figure supplements 9A-B***). The downfield G41 peak (indicator of an inter-molecule hydrogen bond in the homodimer) was absent in the ^15^N-HSQC spectrum of N97A mutant, confirming a global conformation transition from dimer to monomer state. Careful examination of the ^15^N-HSQC NMR spectra of N97A mutant protein samples at concentrations varying from 16 μM to 500 μM revealed clear patterns of concentration dependent shifts (***Figure 5A***). The residues that shifted the most followed a distribution pattern (***Figure 5B***) similar to the chemical shift differences observed between WT and N97A mutant proteins within the GFCC’ face (***Figures 4A, 4B***). This is consistent with the N97A mutant being in rapid monomer-dimer dynamic equilibrium in the fast-to-intermediate exchange regime on the NMR timescale. Fitting the peak trajectories of residues Q44 and A49 provided a rough estimate of over 1 mM for the homodimer dissociation constant K_D_, which agrees well with previous studies (Bonsor et al., 2018).

**Figure 5.**
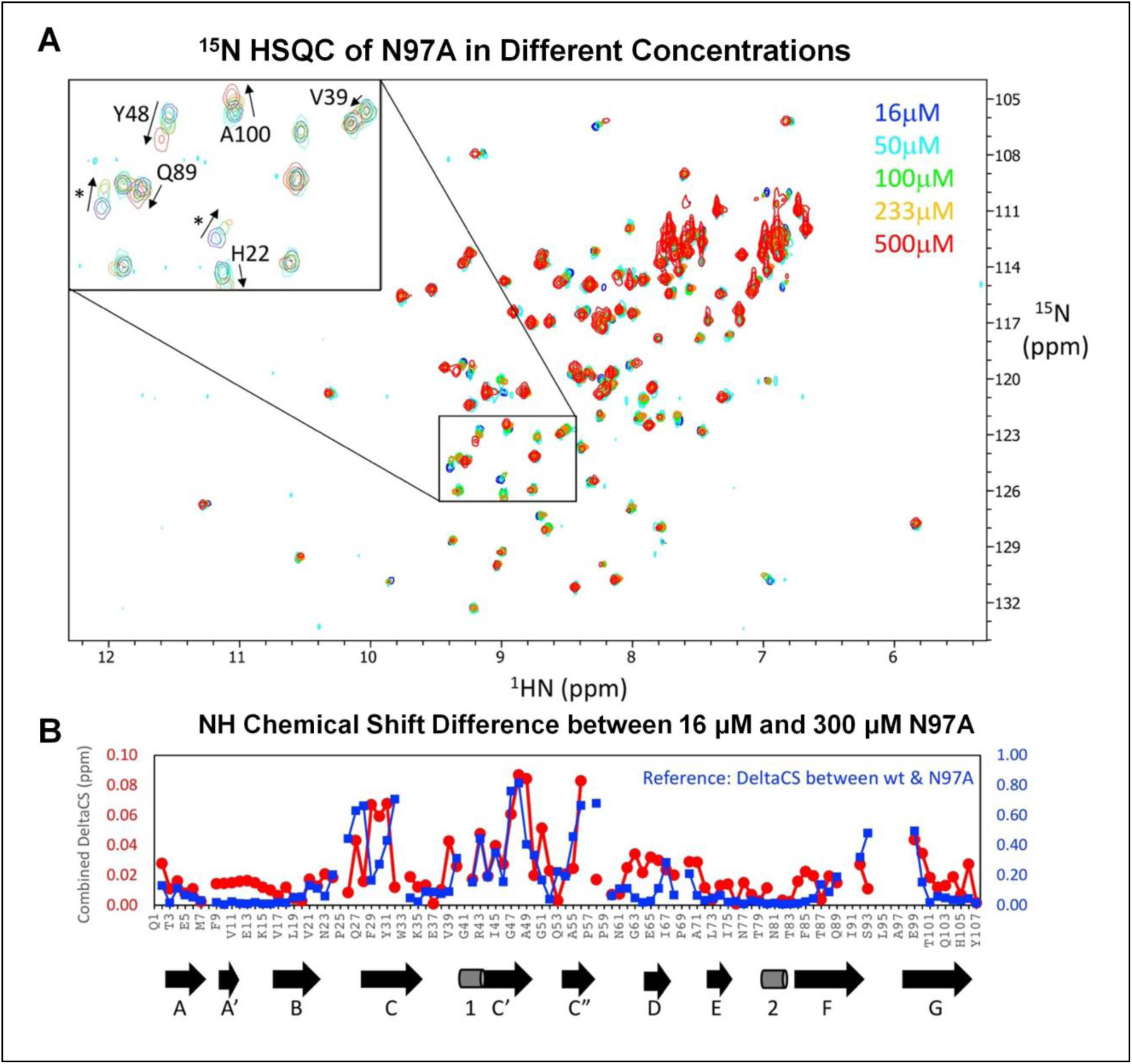
(A) Overlaid ^15^N-HSQC spectra of 16 μM (blue), 50 μM (cyan), 100 μM (green), 233μM (orange) and 500μM (red) N97A mutant hCEACAM1 IgV. The inset shows an enlarged view of a central spectral region containing several assigned residues and unassigned residues (marked with “*”). (B) The relative combined chemical shift changes of assigned ^15^NH peaks between a 16 μM and a 300 μM N97A mutant protein sample (red circles), in comparison with the relative combined chemical shift changes of ^15^NH peaks between the WT and N97A mutant proteins (blue squares). The secondary structure elements from the X-ray structure of N97A protein are depicted below for comparison. **Figure supplement 10** NMR relaxation rates.

In contrast, the ^15^N-HSQC NMR of the hCEACAM1 WT protein samples ranging from 10 μM to 1 mM in concentration did not show concentration dependent shifts (results not shown), consistent with its strong dimer association (K_D_ = 450 nM), and the monomer-dimer equilibrium being in the slow-exchange regime on NMR timescale. The monomer state of the hCEACAM1 WT protein would be expected to appear as minor populations of second conformation peaks in ^15^N-HSQC NMR spectra; however, the peak intensities were too weak to confirm with our NMR sample conditions.

The X-ray crystal structure of the N97A mutant revealed a very similar structural fold compared to the hCEACAM1 WT protein. Although our NMR data largely support the notion that the overall structural fold of N97A in solution is very similar to the X-ray structure model, there was a region of ambiguity around the FG loop. A stretch of nine residues including three residues from the end of the F strand, five residues in the FG loop and one residue at the beginning of the G strand could not be assigned using the conventional NMR sequential connectivity method. On the other hand, there were at least seven NMR peaks in the ^15^N-HSQC spectra with weak or moderate intensities that remained unassigned because of weak NMR cross-correlation peaks. It should be noted that a two-residue fragment was tentatively assigned to K92 and S93 in the FG loop based on it containing the only missing serine residue. It is unlikely that this is due to protein aggregation because there were no improvements when using a triple labeled protein sample that was deuterated to reduce NMR relaxation for large proteins. One explanation is that the FG loop of the N97A monomer was undergoing intermediate rate exchange of multiple conformational states, likely related to the dynamic monomer-dimer equilibrium and/or additional conformational changes.

In addition, ^15^N NMR relaxation studies of hCEACAM1 N97A were carried out by measuring the longitudinal T_1_, transverse T_2_(CPMG) and T_1ρ_ relaxation times (***Figure 5-figure supplement 10A***). The increased R_2_/R_1_ ratios (higher local τ_C_ times) for several residues in the C, C’, C” strands, and potentially the FG loop region, were initially interpreted as reflecting local dynamic motions. However, the differences between the transverse relaxation rates R_2_(CPMG) and R_1ρ_ showed the same pattern, and clearly confirmed its origin to be from NMR chemical shift exchange processes (***Figure 5-figure supplement 10B***). This is because these residues exhibited relatively large ^15^N chemical shift differences, such that the exchange rate from the monomer-dimer equilibrium (at 300μM N97A) falls within the intermediate time scale (on the order of several milliseconds). These results underscore the significant effects that the N97A mutation has on disrupting the extremely stable dimeric form of WT hCEACAM1, and which strongly shifts the dynamic monomer-dimer exchange to a monomeric form.

### 4. Conformation and thermal motion analysis of hCEACAM1 WT IgV and GFCC’ face variants

To investigate the conformational flexibility of the hCEACAM1 IgV at atomic resolution, particularly that associated with the CC’ and FG loops, we first compared the previously published crystal structures of hCEACAM1 WT (PDB code 4QXW, PDB code 2GK2) and hCEACAM1 WT-HopQ complex (PDB code 6AW2) *(Bonsor et al., 2018; Fedarovich et al., 2006; Huang et al., 2015*). Although structural superimposition of a hCEACAM1 molecule from the WT homodimer (PDB code 4QXW) revealed a similar GFCC’ global fold with (RMSD) of 0.635 Å (over 724 atoms) and 0.449 Å (over 598 atoms) for the hCEACAM1 WT structure (PDB code 2GK2) and hCEACAM1-HopQ complex (PDB code 6AW2), respectively, significant conformation differences were observed in the FG loops (***Figure 6-figure supplements 11A, B***). Next, we determined the crystallographic Debye-Waller factor (temperature factor or B factor) of the hCEACAM1 WT homodimer (PDB code 4QXW) and GFCC’ face variant crystal structures described above, expect for the I91A structure, which was determined at too low a resolution (***Table 1***). The B factor can be used to estimate the thermal motion or dynamic mobility and disorder of each atom from the spread of the electron density about the atomic position (*Ringe and Petsko, 1985*). We observed an overall B factor of 23 Å^2^ for a hCEACAM1 WT dimer (PDB code 4QXW) with similar B factor values observed across the GFCC’ face (***Table 1***). In comparison, we observed overall B factors of 28/16/31 Å^2^ for V39A/N97A/E99A, respectively, in the refined crystal structures (***Table 1***) that negatively correlated with their respective calculated melting temperatures (***Figure 1A***) where increased B factor was associated with reduced thermal stability. Although the crystal structure of the N97A mutant showed the lowest B factor value of 16 Å^2^, a closer examination revealed higher B factor profiles of the CC’, EF and FG loop residues (***Figure 6A***).

**Figure 6.**
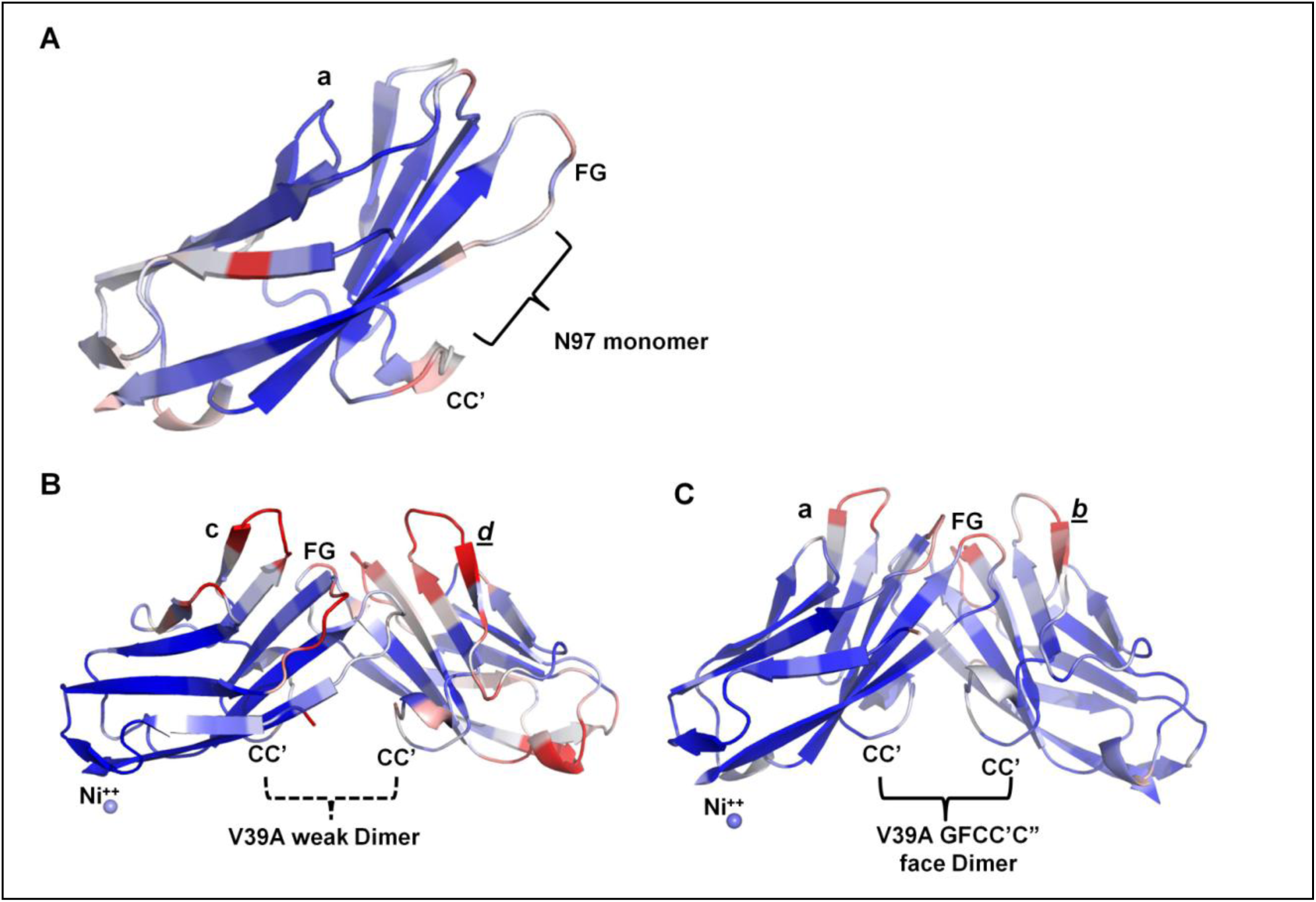
B-factor assignment of N97A and V39A crystal structures. (A) The ribbon diagram of the molecule (a) of N97A crystal structure. The loops, α helices, and β strands are colored based on B-factor range (blue-white-red, where blue minimum=10, red maximum=20). (B) Ribbon diagram of the weak V39A dimer as observed in the V39A crystal structure. The bound Ni^++^, loops, α helices, and β strands are colored based on B-factor range (blue-white-red, where blue minimum=20, red maximum=40). **Figure supplement 11** The conformational flexibility of the hCEACAM1 WT GFCC’ face.

Although packing constraints combined with the time-and space averaging of a crystal structure determination mean that crystalline proteins do not reflect all the characteristics of a protein’s behavior in solution (*Tilton et.al., 1992*), the higher B factor of the CC’ and FG loop relates well with possible multiple conformations states of movement of the CC’ and FG loop residues we observed in our NMR experiments. Following a similar trend, the weaker V39A dimer (molecules c and *d*) showed higher overall B value profiles across the GFCC’ face (***Figure 6B***) compared to the V39A dimer (molecules a and *b*) that resembled a WT dimer (***Figure 6A***). Thus, these conformation and B factor analyses relate well with a dynamic conformational movement of the CC’ and especially FG loop residues we observed in NMR spectroscopy studies of the N97A mutant and support the weakened dimer state of the V39A mutant as a possible transition state as observed in the crystal structure.

## DISCUSSION

We performed biophysical, high resolution structural and NMR studies to determine the basis for CEACAM1 monomer-dimer exchange at an atomic level. Our initial biophysical studies confirmed the previously described dimer-disruptive property of the N97A mutant in solution (*Bonsor et al., 2018*) and demonstrated that the hCEACAM IgV domain exchanges between monomer and dimer forms. Furthermore, introduction of site specific alanine substitutions at the GFCC’ face (V39A, I91A, N97A, E99A) provided for an experimental opportunity to shift the monomer-dimer equilibrium towards each oligomerization species, highlighting the unique thermally stable N97A monomeric variant. Crystal structural analysis of each hCEACAM1 IgV mutant provided static snapshots that included a possible monomeric transition state (V39A) and complete monomeric state (N97A) and NMR studies demonstrated the dynamic properties of the CC’ and FG loops of the N97A mutant in solution.

These studies focus attention on the GFCC’ face and its role in determining the equilibrium between hCEACAM1 monomeric and dimeric species. The weakening of GFCC’ face-mediated dimer association was manifested by disruption of significant CC’ and FG loop residues interactions in the V39A mutant and complete loss of CC’ and FG loop residues interactions in the N97A mutant, suggesting that the GFCC’ face may sequentially ‘unzip’ and ‘rezip’ in transitioning between a dimeric and monomeric state. Consistent with this hypothesis, our NMR studies of the N97A mutant revealed that monomer-dimer exchange involved residues within the GFCC’ face including V39, Y48, Q89 and A100.

It is therefore interesting that we also observed minor interactions between two N97A monomers present in the crystal lattice through the ABED face. Further, the PDB PISA studies revealed that the interfacial interaction involving the ABED loops was associated with a low complexity score, raising the possibility that this face could serve as an additional minor surface that facilitates CEACAM1 oligomerization. Notably an ABED-mediated homodimerization interface has also been suggested in SPR binding studies (*Klaile et al., 2009*) and described in another crystal structure of unglycosylated hCEACAM1 WT IgV (PDB code 2GK2) (*Fedarovich et al., 2006*). Although the ABED surface contains three sites for carbohydrate modification, an attractive possible contribution of the ABED face is to serve as a secondary homo-oligomerization site that achieves relevance following *trans* GFCC’-initiated homo or heterodimerization by propagating surface CEACAM1 clustering and downstream signal activation.

Another contribution of this study was our ability to fully assign the residues within the NMR spectra of WT hCEACAM1 to 100% completion and provide a comprehensive NMR assessment (∼90% assignment) of the N97A mutant monomer. NMR studies confirmed the largely monomeric nature of the N97A mutant in solution (estimated 77% in monomer and 23% in dimer forms at 300 μM protein concentration) but interestingly a tendency to form higher order soluble oligomers over time that could be accommodated by contributions of GFCC’-mediated interactions and additional minor interactions associated with the ABED face. Further, in the case of the N97A mutant, we were able to assign 90% of the backbone amide resonances for the N97A mutant with an inability to assign residues W33, G41 and seven residues in the FG loop region. The unassigned residues, largely in the FG loop that are most affected by the N97A mutant monomer-dimer conformational change, might be a consequence of exceptionally large ^15^NH chemical shift differences leading to complete exchange line-broadening resulting in extremely weak and/or missing resonance peaks. However, the ^15^N-HSQC spectrum of the 16 μM N97A mutant sample, where only 2% of the proteins were estimated to be dimeric, did not show newly emerged peaks (***Figure 5A***). In addition, although some of the unassigned seven ^15^NH resonance peaks appeared to have stronger intensities at 16 μM compared to higher protein concentrations, they did not show significantly larger chemical shift changes with increasing N97A protein concentrations. Perhaps these cases are more complicated and possibly involve additional conformational changes besides the monomer-dimer equilibrium for these FG loop residues at the GFCC’ face. The higher conformational flexibility and B factor of FG loops residues as described before for WT (PDB code 4QXW, 2GK2) and the hCEACAM1-HopQ complex structure (PDB code 6AW2) (***Figure 6A,B, Figure 6-figure supplements 11A, B)*** support this local structural malleability which could facilitate hCEACAM1 IgV homodimer formation as well as the formation of complexes with many other proteins, including TIM-3 *(Gandhi et al., 2018; Huang et al., 2016; Huang et al., 2015), HopQ (Bonsor et al., 2018)* and other microbial pathogens *(Kim et al., 2019*). Although the residues that exhibited the most spectral change differences between WT and N97A mutant protein were in the same region as the N97A ^15^NH peaks that exhibited the largest shifting caused by changes in N97A monomer-to-dimer equilibrium (***Figure 5B***), the magnitude and direction of the ^15^N-HSQC spectral changes do not match exactly. It also does not appear that the ^15^N-HSQC spectrum of N97A mutant at even higher concentrations (i.e. sample with close to 100% dimer form) will replicate that of the hCEACAM1 WT IgV. Therefore, it is likely that the dimer form of N97A as shown in equilibrium with the monomer could be a transition state dimer akin to what was observed for the V39A mutant. The local structural malleability of FG loop residues is also well supported by thermal motion analysis of the N97A crystal structure wherein we observed higher a B-factor at these locations. Thus, these NMR results support the unique role of the N97 residue in determining the hCEACAM1 monomer-dimer equilibrium and the impact of the V39 hydrophobic interactions on hCEACAM1 homodimerization.

Further, our high resolution structural and NMR studies are consistent with a model wherein the GFCC’ loops of hCEACAM1 represent the primary face involved in homodimer formation (***Figure 7***). When this GFCC’ face is abrogated, possibly first through disruption of CC’ loop interactions (as observed in the V39A weak dimer GFCC’ face crystal structure) or when hCEACAM1 is a monomer (as observed in the N97A crystal structure) in the *cis* state, the higher thermal motion and dynamic conformation state especially associated with the CC’ and FG loops contributes to monomeric hCEACAM1 homophilic behavior or facilitates heterophilic interactions with its other ligands in *cis* or *trans* through the GFCC’ face of hCEACAM1 (***Figure 7***). The importance of the GFCC’ face is also highlighted by the high degree of genetic polymorphisms that exist there, as previously described (*Huang et al., 2015*), suggesting that a genetic propensity and/or other factors as discussed below may be involved in regulating these processes. In the event of homophillic interactions by monomeric hCEACAM1 with a neighboring hCEACAM1 monomer, the hCEACAM1 GFCC’-mediated homodimer is formed and becomes more thermally stable (***Figure 7***). The GFCC’ face stabilized homodimer could subsequently participate in higher order oligomer formation, which we observed in our NMR studies, possibly through minor interactions mediated though the ABED face as observed in our N97A crystal structure or another crystal structure of hCEACAM1 (PDB code 2GK2) (*Fedarovich et al., 2006)*.

**Figure 7.**
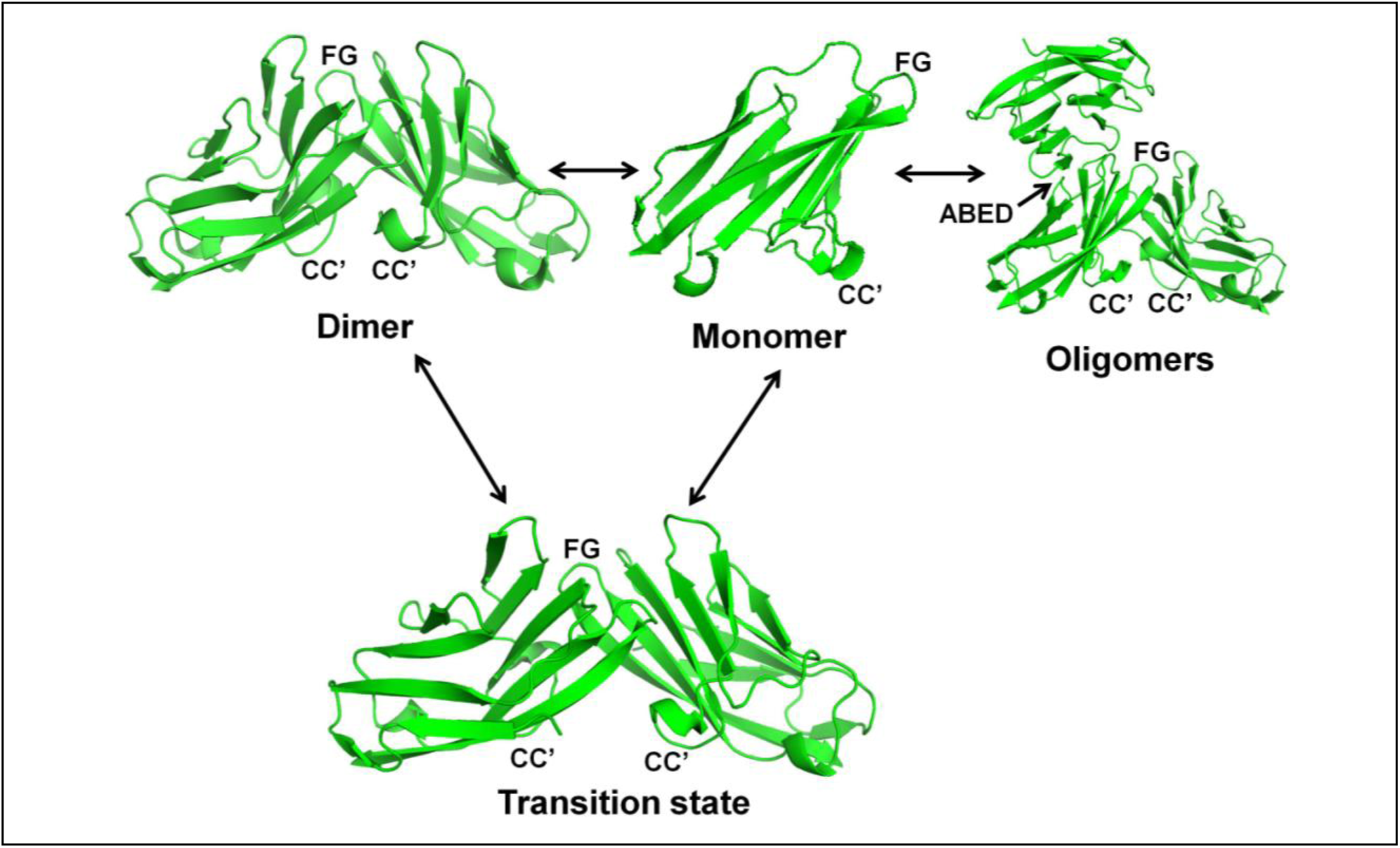
Monomer, dimer, transition and oligomeric states of hCEACAM1 IgV as observed in various crystal structures. The ribbon diagram of all states of hCEACAM1 are shown with labeled CC’ and FG loops. The dimer state of hCEACAM1 was observed in WT crystal structure (PDB code 4QXW) and mediates dimer formation through GFCC’ face. The weak V39A dimer as observed in the crystal structure and described in *Figure 3B* results from weaker hydrogen bonded and hydrophobic interactions at the GFCC’ face and mimic transition state. The N97A mutation results in the complete abrogation of GFCC’ face dimer as described in the *Figures 3C and 3D.* The monomeric state of N97A mutant as observed in the crystal structure is further confirmed by NMR studies. Further, we observed tendency to form higher order soluble oligomers over time in our NMR studies which could possibly occur through association of GFCC’ face dimer with other hCEACAM1 molecules thorough ABED face as observed in N97A crystal structure whereas molecules (a) and (*b*) showed minor point of contact through ABED face. Similar ABED face association was also observed in previously reported structure of hCEACAM1 IgV (PDB code 2GK2).

This would enable interactions between CEACAM1 cytoplasmic tails that facilitate CEACAM1 signaling associated with its inhibitory functions through association with *Src*-homology domain containing phosphatases. Indeed, many functional studies by others have also demonstrated that the propensity of CEACAM1 to form higher order oligomers may be initiated by initial formation of monomers through transmembrane or cytoplasmic tail interactions with calcium followed by GFCC’ face interactions (*Klaile et al., 2009, Patel et al., 2013*). Interestingly, up to 50 percent of CEACAM1 on the cell surface of CEACAM1 transfected cells has been predicted to be in a monomeric state with the remainder existing as homodimers consistent with a monomer-dimer equilibrium in the physiologic function of this important cell surface protein. Thus, our structural model extends our understanding of the hCEACAM1 monomer-dimer equilibrium and provides a structural rationale for oligomerization-mediated activities.

Our findings also help to understand how the dynamic nature of the hCEACAM1 GFCC’ face facilitates its binding with various other host ligands such as hTIM-3 and CEACAM5 along their GFCC’ faces, the proteins associated with numerous bacterial pathogens such as HopQ of *Helicobacter pylori*, Opa proteins of *Neisseriae sp.* and Afa/Dr adhesins of *E. coli* and their potential therapeutic blockade with peptides or monoclonal antibodies (***Figure 7-figure supplement 12***). hTIM-3 in particular is an important immunoregulatory protein that possesses an N-terminal IgV domain with very high structural similarity (main chain RMSD of 1.31 Å) to the hCEACAM1 IgV domain with. We recently solved the hTIM-3 IgV (PDB code 6DHB) crystal structure in association with bound calcium and reported the first NMR study of a TIM family member (BMRB ID 27525) (*Gandhi et al., 2018)*. Further, using various cellular, biochemical and biophysical methods (*Gandhi et al., 2018; Haidar et al., 2019; Y.-H. Huang et al., 2016; Sabatos-Peyton et al., 2018; Zhang et al., 2020)*, we and others have demonstrated a conserved role of the GFCC’ faces of hCEACAM1 and hTIM-3 in the heterodimerization between these two proteins (K_D_ of ∼ 2-3 μM) (***Figure 7-figure supplement 12***). Despite these findings, a recent paper (*De Sousa Linhares et al., 2020*) suggested a lack of appreciable binding of hCEACAM1 by hTIM-3. This is surprising and likely due to a number of factors, including use of incompletely characterized Fc fusion proteins that do not take into account the monomer-dimer equilibrium or structural state of the proteins used or consideration of the relative affinities of the CEACAM1 homodimer (∼450 nM) and CEACAM-TIM-3 heterodimer (∼2-3 μM). In the ELISA studies for example a single concentration of immunoglobulin fusion proteins was used in the nanomolar range without the control of the calcium level, which naturally shifts the interaction towards detection of higher affinity hCEACAM1 homophillic binding rather than lower affinity hCEACAM1-hTIM-3 interactions. Further, there is an absence of titration experiments in the micromolar range to probe the binding and there are confounding results that show that the strongest ligand binding to the hCEACAM1-Ig used was with galectin-9 in the absence of galectin-9 binding to hTIM-3. Importantly, galectin-9 has never been described as a ligand for hCEACAM1 and numerous groups have unambiguously identified an interaction between hTIM-3 and galectin-9 *(Cao et al., 2007; van de Weyer et al., 2006; Zhang et al., 2020; Zhu et al., 2005*), raising important concerns about the interpretation of these and other results contained therein (*De Sousa Linhares et al., 2020)*.

The homo-oligomeric and hetero-oligomeric properties of CEACAM1, especially with regards to TIM-3 and microbial ligands, carries significant therapeutic potential making our understanding of the CEACAM1 monomer to dimer transition and associated receptivity of the CEACAM1 monomer of great importance. A number of groups (*Gandhi et al., 2018; Haidar et al., 2019; Sabatos-Peyton et al., 2018; Zhang et al., 2020)* have recently observed that selective targeting of the GFCC’ faces of either hCEACAM1 (e.g. with the 5F4 monoclonal antibody or hTIM-3 peptides) or hTIM-3 (e.g. with polyclonal antibodies or monoclonal antibodies including 2E2 or M6903) can disrupt formation of hCEACAM1 homodimers and complexes with hTIM-3, respectively, using therapeutic agents that exceed the natural homodimeric and heterodimeric affinities (***Figure 7-figure supplement 12***). These results are consistent with the importance of amino acid residues such as E62 and D120 within the hTIM-3 GFCC’ face that determine binding to hCEACAM1 as defined by site-directed mutagenesis (*Huang et al., 2015*). The hTIM-3 bound crystal structure of the M6903 anti-TIM3 monoclonal antibody that blocks hCEACAM1-hTIM-3 interactions revealed the antibody Fab binds with TIM-3 residues E62 and D120 at atomic level resolution (*Zhang et al., 2020*). Similarly, hydrogen deuterium exchange mass spectrometry (HDxMS) studies revealed that the anti-hTIM-3 monoclonal antibody 2E2 binds the GFCC’ face of hTIM-3 with blockade of hCEACAM1 binding in transfected cells based upon flow cytometry (*Sabatos-Peyton et al., 2018)*. Importantly, amino acid residues such N42, R43, Q44, G47, and Q89 of hCEACAM1 that mediate hTIM-3 binding as defined by site-directed mutagenesis studies are also located within the GFCC’ face and involved in the formation of hCEACAM1 homodimers (***Figure 7-figure supplement 12***). This highlights the strong competition between formation of high affinity homodimers and lower affinity heterodimers and the critical need to take the monomer-dimer equilibrium into account in considering and designing experimental conditions to detect hCEACAM1 interactions with its various ligands, a note of caution given the recent studies of others (*De Sousa Linhares et al., 2020)*.

Consistent with our findings, many recent biophysical studies, including the crystal structure of a hCEACAM1-HopQ complex and small angle x-ray scatting studies, support the importance of the GFCC’ face and monomer-dimer equilibrium in the binding of hCEACAM1 to its various ligands (***Figure 7-figure supplement 12***). In the case of HopQ, its higher affinity for hCEACAM1 (K_D_ ∼23-279 nM) relative to that associated with hCEACAM1 homodimerization (K_D_ ∼450 nM) resulted in successful competition for binding to hCEACAM1 GFCC’ face of another monomer. Furthermore, the crystal structure that defines this ability of HopQ to achieve the formation of a high affinity heterodimer is through its competence in interacting with the V39, I91 and N97 residues of hCEACAM1 (***Figure 7-figure supplement 12***). Similarly, hCEACAM1 GFCC’ face residues F29, Q44, I9I and E99 have also been shown to be critical to the binding of OPA proteins and Afa/Dr adhesins (***Figure 7-figure supplement 12***). Thus, these recent findings further extend the role of GFCC’ face residues not only in the formation of hCEACAM1 homodimers, but also in the interactions with many different heterophilic ligands including TIM-3, CEACAM5, HopQ, Opa proteins and Afa/Dr adhesins in immune-regulation and immune-evasion that are exploited by neoplastic cells and microbial pathogens (***Figure 7-figure supplement 12***).

In summary, our biophysical and structural studies by crystallography and in solution by NMR support a model wherein the GFCC’ face is highly dynamic and seeks thermal and energetic stability through formation of dimers (either homodimers or heterodimers) that lock in a structurally favorable state. As such, CEACAM1 prefers to be in a dimeric state that specifically stabilizes the CC’ and FG loops, making the GFCC’ face the major interaction site for formation of homodimers and heterodimers. A caveat of these studies is that they were performed with unglycosylated proteins that may affect the ABED face; however, studies reporting a role of glycosylation in disrupting homodimerization have been retracted (*Zhuo et al., 2020, Zhuo et al., 2016*). That said, given the location of the carbohydrate side-chain modifications of CEACAM1 along the ABED face, the mutational analyses performed here and its implications still have substantial physiologic merit. In addition to understanding the structural mechanisms that underlie the formation of CEACAM1 monomers that are amenable to interactions with another CEACAM1 molecule or its potential heterophilic partners, we are hopeful that our high-resolution structural studies and hCEACAM1 monomer-dimer model may also be useful in considering therapeutic targeting of hCEACAM1 interactions with its various ligands.

## Materials and methods

### Protein Expression and Purification

The hCEACAM1 WT IgV and mutant (V39A, I91A, N97A, E99A) proteins were expressed and purified using our published protocols (*Huang et al., 2016*).

### Differential Scanning Fluorimetry

Differential scanning fluorimetry was performed using a QuantStudio 6 (Life Technologies) RT-PCR instrument with the excitation and emission wavelengths set to 587 and 607 nm, respectively. Assay buffer was 10 mM HEPES pH 7.4, 150 mM NaCl. For thermal stability measurements, the temperature scan rate was fixed at 1 °C/min. Protein concentration was uniform at 25 μM among the hCEACAM1 WT and mutant samples and SYPRO orange (Invitrogen) concentration was consistent at 5x. The temperature range spanned 20 °C to 95 °C. Data collection was performed by Quant Studio Real-Time PCR Software (Life Technologies) on triplicate samples and analyzed by Protein Thermal Shift Software v1.4 (ThermoFisher). Melting point temperature (*T*_m_) was calculated for each protein samples through computation of a temperature derivative for each respective melting curve that was then processed with a peak fitting algorithm, applying a sigmoidal baseline and fitting the peak to determine the *T*_m_ and its standard error.

### Size-Exclusion Chromatography with Multi-Angle Light Scattering

Purified samples of WT and mutant (V39A, I91A, N97A, E99A) hCEACAM1 IgV were evaluated for size and monodispersity by analytical size exclusion chromatography and multi-angle light scattering (SEC-MALS). Samples of hCEACAM1 at 100 μM were injected onto a TSK-gel Bioassist G4SWxl (Tosoh) SEC column, equilibrated with 10 mM HEPES pH 7.4, 150 mM. The SEC column was coupled to a static 18-angle light scattering detector (DAWN HELEOS-II) and a refractive index detector (Optilab T-rEX) (Wyatt Technology, Goleta, CA). Data were collected at a flow rate of 0.5 mL/min. Retention time, molecular weight and polydispersity index (PDI) were calculated in ASTRA (Wyatt).

### Crystallization, Data Collection, and Structure Determination

Purified mutant (V39A, I91A, N97A, E99A) proteins were concentrated to 10 mg/ml (exceeding 800 μM concentration) and preliminary crystallization screens were performed using Index HT (Hampton Research) and PEGRx (Hampton Research) at 4 °C and room temperature. The conditions with promising hits were later optimized with an additive screen (Hampton Research) and x-ray diffraction quality crystals were obtained under crystallization conditions of 0.005 M Cobalt(II) chloride hexahydrate, 0.005 M Nickel(II) chloride hexahydrate, 0.005 M Cadmium chloride hydrate, 0.005 M Magnesium chloride hexahydrate with 12% w/v Polyethylene glycol 3,350 in 0.1 M HEPES pH 7.5 buffer (V39A mutant condition), 54% Tascimate with 0.5 % n-Octyl-β-D-glucoside pH 8.0 (I91A mutant condition), 6% Tascimate with 25% polyethylene glycol 400 in 0.1 M MES monohydrate pH 6.0 (N97A mutant condition), and 41% Tascimate with 0.5 % n-Octyl-β-D-glucoside pH 8.0 (E99A mutant condition), respectively. For x-ray data collection, crystals were cryoprotected in mother liquor of crystallization condition with approximate concentration of 12% glycerol and 7% ethylene glycol. X-ray data for V39A, I91A, and E99A mutant protein crystals were collected at the 21-ID-F/G, LS-CAT beamlines at the Advanced Photon Source (APS; Argonne, IL, USA) and at the 17-ID beamlines, National Synchrotron Light Source-II (NSLS-II, Upton, NY, USA) for the N97A mutant protein crystals. The diffraction data for each of the mutant (V39A, I91A, N97A, E99A) protein crystals were processed with iMosflm and the CCP4 suite of software *(Battye et al., 2011; Winn et al., 2011*), HKL2000 (*Otwinowski & Minor, 1997*), and FastDP (*Winter & McAuley, 2011*) in-house and at the beamlines (APS, Argonne, IL; NSLS-II, Upton, NY). The structure of the each mutant was determined by molecular replacement with MolRep (*Winn et al., 2011*) using a polyalanine model of our hCEACAM1 WT crystal structure (PDB code 4QXW) and many rounds of structure refinement were done with simultaneous model building using Refmac (*Murshudov, Vagin, & Dodson, 1997)* and COOT (*Emsley et al., 2010*). The Fo-Fc map at 3.0 σ level (derived from the initial model) showed significant positive Fo-Fc map density where residue A39 in the V39A refinement model, residues A91 in the I91A refinement model, residue A97 in the N97 refinement model, and residues A99 in the E99A refinement model were not fitted, respectively (***figure supplements 13A-D***).

The crystallographic twining and the significant metal electron density near the FG loop were observed in the V39A protein during the data processing and the structure refinement. The V391 mutant protein crystallization condition from Index HT screen (Hampton Research) has 0.005 M Cobalt(II) chloride hexahydrate, 0.005 M Nickel(II) chloride hexahydrate, 0.005 M Cadmium chloride hydrate, and 0.005 M Magnesium chloride hexahydrate. Using intensity based twin refinement in Refmac and looking at the interacting residues H105 and V106 around the observed metal density, we resolved the final crystal structure with bound Ni**^++^** (Nickel) for the molecule (a) or (c) and their two symmetry mates 0100-100 (S1) and 02000000 (S2) with the strategy that showed optimal R/Rfree and revealed hexadentate interactions of Ni**^++^** with three His105 sidechains and three carbonyl groups of Val106 residues (***Figure 3-figure supplements 5B-C***). The final crystal structure of the V39A mutant was determined in the P3 space group with Rwork/Rfree of 14.51%/19.39 % at 1.9Å resolution. For other mutants, the crystal structure of the I91A mutant was solved in the P4212 space group with Rwork/Rfree of 22.1%/25.8 at 3.1Å resolution, that of the N97A mutant in the C2221 space group with Rwork/Rfree of 19.18%/23.99% at 1.76Å resolution, and the E99A mutant in the P4212 space group with Rwork/Rfree of 18.94%/22.32 at 1.9Å resolution, respectively. In order to further validate observed 2Fo-Fc electron density of each mutant at the mutation site (***figure supplements 13E-H)***, the final refined model of each mutant was changed to the original residue as present in the hCEACAM1 IgV WT (valine for the V39A refined model, isoleucine for the I91A refined model, asparagine for the N97A refined model, and glutamic acid for the E99A refined model) and an additional cycle of refinement was performed. The significant negative Fo-Fc map at 3.0 σ level for each mutant at the reverse mutation site was observed and validated structures of V39A, I91A, N97A, and E99A mutants (***Figure supplements 13I-L)***.

The atomic coordinates and structure factors were deposited with RCSB accession code 6XNW (V39A), 6XNT (I91A), 6XNO (E99A), 6XO1 (N97A), respectively. The X-ray data and structure refinement statistics for the V39A, I91A, N97A, E99A mutant crystal structures are shown in Table 1. All the figures, B factor calculation and conformational mapping were carried out using PyMOL (DeLano Scientific) and sequence alignments of hCEACAM1 were done using Clustal Omega (*Sievers et al., 2011*).

**Table 1:** Crystal information, data collection and refinement parameters

|  | V39A (PDB<br>code 6XNW) | I91A (PDB<br>code 6XNT) | E99A (PDB<br>code 6XNO) | N97A (PDB<br>code 6XO1) |
| --- | --- | --- | --- | --- |
| **Data collection statistics** |  |  |  |  |
| **Space Group** | P 3 | P 4212 | P 4212 | C 2221 |
| Cell constants<br>a (Å), b (Å), c (Å),<br>$\alpha^\circ$, $\beta^\circ$, $\gamma^\circ$ | 91.44, 91.44, 64.41<br>90.00 90.00 120.00 | 102.11, 102.11,<br>61.02, 90.00 90.00<br>90.00 | 106.82, 106.82,<br>62.23, 90.00<br>90.00 90.00 | 55.93,56.88,124.54<br>90.00 90.00 120.00 |
| Resolution (Å)* | 39.59-1.90 ( 1.94-1.90) | 72.21-3.1 (3.31-3.1) | 75.5-1.9(1.94-1.90) | 28.760-1.76 ( 1.8-1.76) |
| No. of measurements* | 366913 (23237) | 73320 ( 12990) | 393727(24522) | 231087 (6712) |
| Unique reflections* | 47459( 3048) | 6265 ( 1107) | 28973 (1843) | 19429 (1087) |
| I/sigma I* | 7.8 (1.4) | 9.4 ( 3.8) | 14.3 (2.3) | 19.5 (2.0) |
| Completeness (%)* | 100 (100) | 100 (100) | 100 (100) | 96.1(73.2) |
| Redundancy* | 7.7 (7.6) | 11.7 (11.7) | 13.6 (13.3) | 11.9 (6.2) |
| $R_{merge}$ (%)<sup>a,*</sup> | 18.8 (155.1) | 24.6( 75.4) | 11.9 (140.3) | 8.6 ( 833) |
| $CC_{1/2}$ * | 0.992 (0.304) | 0.963 ( 0.894) | 0.998 ( 0.769) | 0.998 (0.794) |
| **Structure refinement** |  |  |  |  |
| $R_{work}$ (%)<sup>b,*</sup> | 14.51 | 22.1 | 18.94 | 19.18 |
| $R_{free}$ (%)<sup>c,*</sup> | 19.39 | 25.8 | 22.32 | 23.99 |
| No. atoms |  |  |  |  |
| Protein/Water | 3373 | 1720 | 1814 | 1870 |
| R.m.s.deviation |  |  |  |  |
| Bond-lengths (Å) | 0.031 | 0.014 | 0.021 | 0.019 |
| Bond-angles (°) | 2.93 | 1..85 | 2.15 | 1.916 |
| **Ramachandran** |  |  |  |  |
| Favored regions (%) | 94.05 | 94.29 | 96.67 | 97.14 |
| Allowed regions (%) | 4.76 | 4.76 | 2.38 | 2.86 |
| Disallowed (%) | 1.19 | 0.95 | 0.95 | 0 |
| **B-factors, All atoms (Å<sup>2</sup>)** | 28 | 42.0 | 31.0 | 16.0 |

<sup>a</sup> $R_{merge} = \sum |I - \langle I \rangle| / \sum \langle I \rangle$ , where $I$ is the observed intensity and $\langle I \rangle$ is the weighted mean of the reflection intensity.
<sup>b</sup> $R_{work} = \sum ||F_o| - F_c| / \sum |F_o|$ , where $F_o$ and $F_c$ are the observed and calculated structure factor amplitudes, respectively.
<sup>c</sup> $R_{free}$ is the crystallographic $R_{work}$ calculated with 5 % of the data that were excluded from the structure refinement.
\* Values in the parentheses are for highest resolution shell.

### Nuclear Magnetic Resonance Studies

^15^N/^13^C double-labeled WT and N97A hCEACAM1 IgV proteins were expressed from *E. coli* in M9 minimal medium containing ^15^NH_4_Cl and ^13^C-gluocse as the sole nitrogen and carbon sources and purified as described above. ^15^N/^13^C/^2^H triple-labeled WT and hCEACAM1 IgV proteins were expressed similarly except in D_2_O instead of H_2_O. Non-uniformly sampled (NUS) triple resonance experiments using ^15^N/^13^C/^2^H-hCEACAM1 IgV (0.2mM) in 10mM HEPES, 50mM NaCl, pH 7.0 with 10% D_2_O, were performed at 25^°^C on a 700MHz Agilent DD2 spectrometer equipped with a cryogenic probe. The data were processed using NMRPipe (*Delaglio et al., 1995*) and Iterative Soft Thresholding reconstruction approach (istHMS) (*Hyberts et al.*, *2012)* and analyzed by CARA (*Keller, R 2004*). Backbone assignment experiments for N97A mutant hCEACAM1 IgV were performed with a 0.3mM ^15^N/^13^C double-labeled protein sample under the same condition using the same methods as described above. The assigned NMR chemical shifts have been deposited in the Biological Magnetic Resonance Bank database (BMRB ID 50368 for hCEACAM1 IgV WT dimer and BMRB ID 50366 for N97A mutant hCEACAM1 IgV monomer. Secondary structure predications based on assigned chemical shifts (H, HN, CO, CA and CB) were obtained using the TALOS-N software (*Shen & Bax, 2013*)

The 1D ^15^N TRACT (Lee 2006) experiments were carried out using ^15^N/^13^C labeled N97A mutant hCEACAM1 IgV protein samples at 25°C on a 500MHz Bruker Avance III HD spectrometer. Series of relaxation delays (in increments of 40ms or 50 ms) up to 200ms or 250ms were used for pro TROSY measurements; and delays (in increments of 15ms or 20ms) up to 60ms or 100ms were used for anti TROSY measurements. The 1D spectral region between 9.5ppm and 8.7ppm were integrated to extract the pro and anti TROSY relaxation rates. Data were processed and analyzed using the Bruker Topspin program.

NMR relaxation experiments were carried out using a 0.3mM ^15^N/^13^C double-labeled N97A mutant hCEACAM1 IgV protein sample at 25°C on a 700MHz Agilent DD2 spectrometer equipped with a cryogenic probe. The T_1_ relaxation times were determined using antiphase inversion recovery delays of 10ms, 250ms, 500ms, 750ms and 1s. The T_2_ relaxation times were determined using Carr-Pursell-Meiboom-Gill (CPMG) pulse train with τ value of 625μs, and delays of 10ms, 30ms, 50ms, 90ms and 150ms. The T_1rho_ relaxation times were determined with spin-locking field strength of 1.5kHz, and delays of 10ms, 30ms, 50ms, 90ms and 130ms. Data were processed using NMRPipe and analyzed by CARA and Microsoft Excel programs. The errors of the relaxation times were estimated from fitting routines. The average value of R_2_/R_1_ (=T_1_/T_2_) relaxation rate ratio is 9.4. This corresponds to a molecular rotational correlation time of 7.9ns 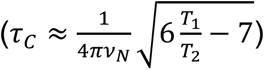, in agreement with the TRACT measurement results after taking into account the homodimer population. The NMR chemical exchange process for some of the N97A residues were not sufficiently suppressed by the T_2_ CPMG pulse train (with τ = 625μs), versus the more efficient rotation frame T_1ρ_ spin-locking, resulting in artificially higher R_2_/R_1_ and R_2_/R_1ρ_ ratios.

## PDB PISA Validation

The PDB PISA (Proteins, Interfaces, Structures and Assemblies) validation was done using the final refined atomic coordinate file of each mutant (V39A, I91A, N97A, E99A) and complex significance scores (CSS) with residue level interactions details were determined (*Krissinel & Henrick, 2007*). The hydrogen bond interactions and CSS scores as observed in the crystal structures of the V39A, I91A, N97A, E99A mutants are shown in supplements table 2 and table3, respectively.

## Acknowledgements

The authors thank all the beamline scientists and the support staff of LS-CAT beamlines 21-ID-F and21-ID-G (APS; Argonne, Illinois, USA) and NSLS-II 17-ID-1 and 17-ID-2 beamlines (Upton, NY, USA) for x-ray data collection, and East Quad NMR (EQNMR, Boston, MA, USA) and Dana-Farber Cancer Institute NMR (DFCI Boston, MA, USA) facilities for NMR data collection. We are thankful to M. Pyzik and A. Riar for helpful literature insights and discussions. This work was supported by the NIH Grant 5R01DK051362-21 and the High Pointe Foundation to R.S.B., and 5P01AI073748-09 to V.K.K. R.S.B. has several issued and pending patents describing potential therapeutic strategies for inhibiting CEACAM1.

## Author Contributions

A.K.G., W.M.K. and Z.-Y.J.S. performed biophysical characterization, x-ray crystallography and NMR experiments. W.M.K. performed DSF and SEC-MALS experiments. A.K.G. performed x-ray crystallization, structure determination and conformational analysis of hCEACAM1 mutants. Z.-Y.J.S. and GW planned NMR experiments. Z.-Y.J.S. carried out NMR experiments and assigned NMR spectra of the hCEACAM1 IgV dimer and N97A monomer. Y.-H.H., and Y.K. designed expression constructs and performed similarity analyses of hCEACAM family members. D.A.B. and E.J.S. established purifications and refolding protocol and conceptualization of conformational analysis. G.P. provided structural expertise in crystallization, refinement, B factor analysis and structural comparison. R.S.B. and V.K.K. established this hCEACAM1 project, and together with A.K.G., G.P., W.M.K., Z.-Y.J.S., Y.-H.H. and Y.K. wrote the manuscript. R.S.B. and A.K.G are the senior author of this paper.

## Supplement figures

**Figure supplement 1.**
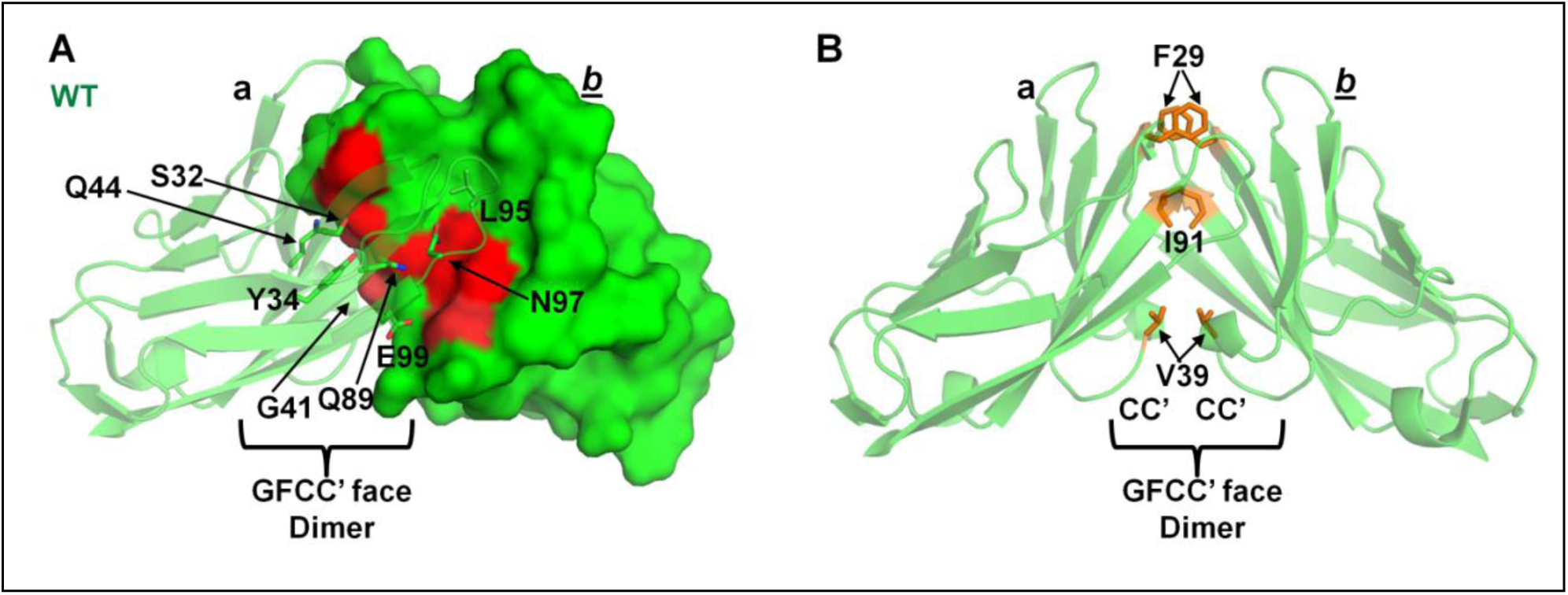
Crystal structure of the IgV of hCEACAM1 homodimer (PDB code 4QXW). (A) Ribbon (molecule a) and surface diagram (molecule *b*) of the hCEACAM IgV crystal structure in green. The residues S32, Y34, G41, Q44, Q89, L95, N97, and E99 that mediates hydrogen bonded interactions to form GFCC’ face dimer are shown by stick and surface (bright and light red) representation. The molecule (*b*) residues which make multiple interactions across the GFCC’ face are shown by bright red surface representation. (B) The hCEACAM1 residues F29, I91, and V39 of molecule (a) and (*b*) that mediate strong hydrophobic interactions in the formation of GFCC’ face dimer are shown by orange stick representation. The CC’ loops are labeled.

**Figure supplement 2.**
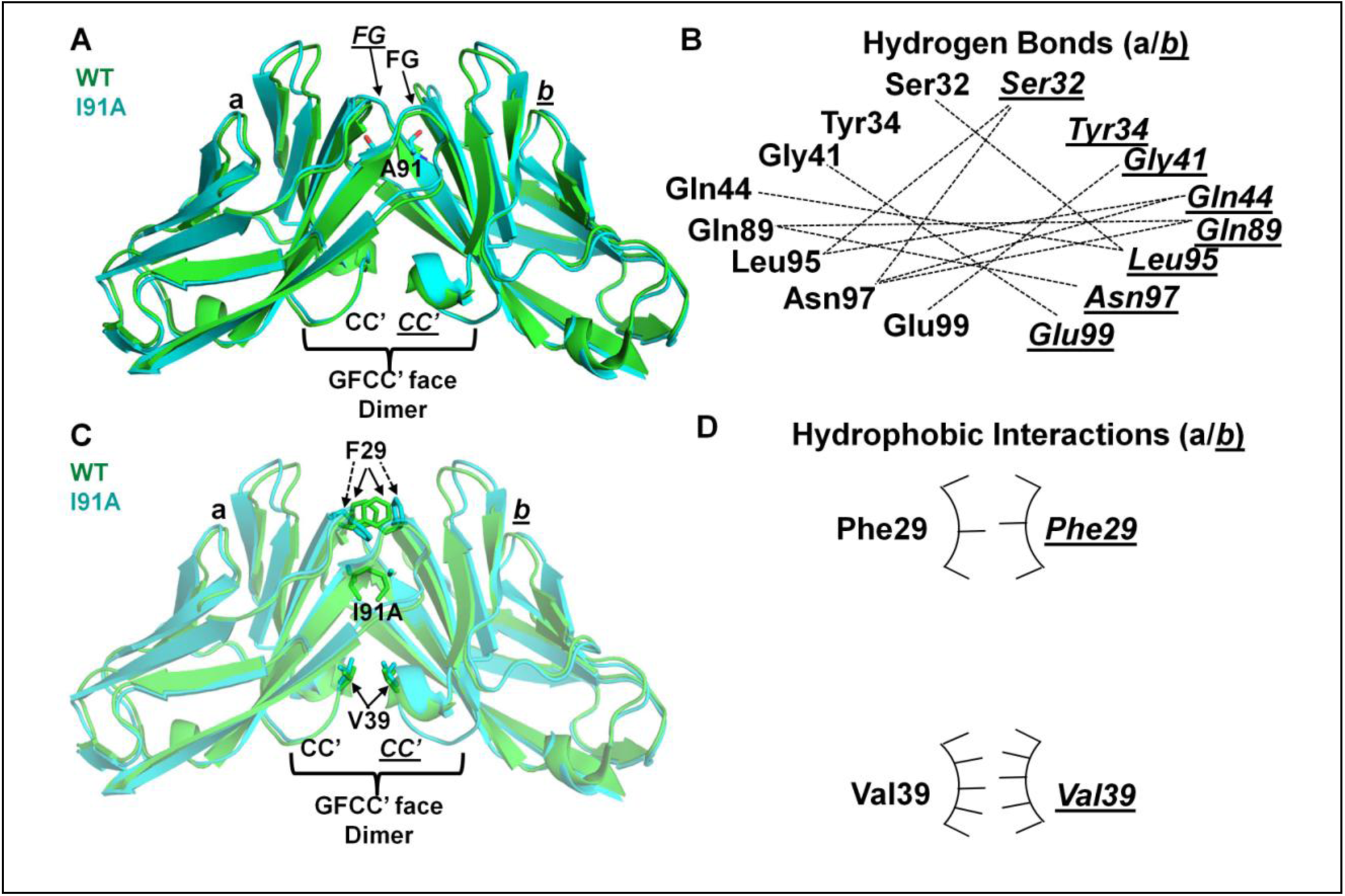
Structural comparison of the I91A mutant and hCEACAM1 WT IgV (PDB code 4QXW) crystal structures. Ribbon diagram of the I91A mutant (cyan) and WT (green) IgV molecules (a) and (*b*) superimposed on each other and GFCC’C’ face dimer is shown. The CC’ and FG loops of WT and I91A mutant are labeled in bold and Italics underlined, respectively. The A91 residues as observed in the I91A crystal structure is shown by stick representation. (B) The hydrogen bonded interactions between the residues described in the Figure 2B are shown by dashed lines, where interactions of the residues (labeled bold) of the molecule (a) and residues (italics underlined) of the molecule (*b*) mediates formation of GFCC’ face dimer. (C) The superimposition of the I91A mutant (cyan) and WT (green) IgV molecules (a) and (*b*), where residues F29 and V39 involved in hydrophobic interactions are shown by stick representations. The F29 residues (cyan, dashed arrows) of I91A mutant make weaker hydrophobic interactions compared to WT F29 residues (green, solid arrows). The site of I91A mutation on the superimposed I91A mutant and WT structure is shown by stick representation. The CC and CC’ loops involved in the formation of GFCC’C’ face I91A and WT dimer are labeled. (D) The weaker hydrophobic interactions as observed in the I19A crystal structure shown by arc/stick representation, where F29 and V39 residues are labeled in bold and italics underlined for molecules (a) and (*b*), respectively. The weaker hydrophobic interactions between two F29 residues of I91A mutant are shown by less numbers of pointers on the hydrophobic arc.

**Figure supplement 3.**
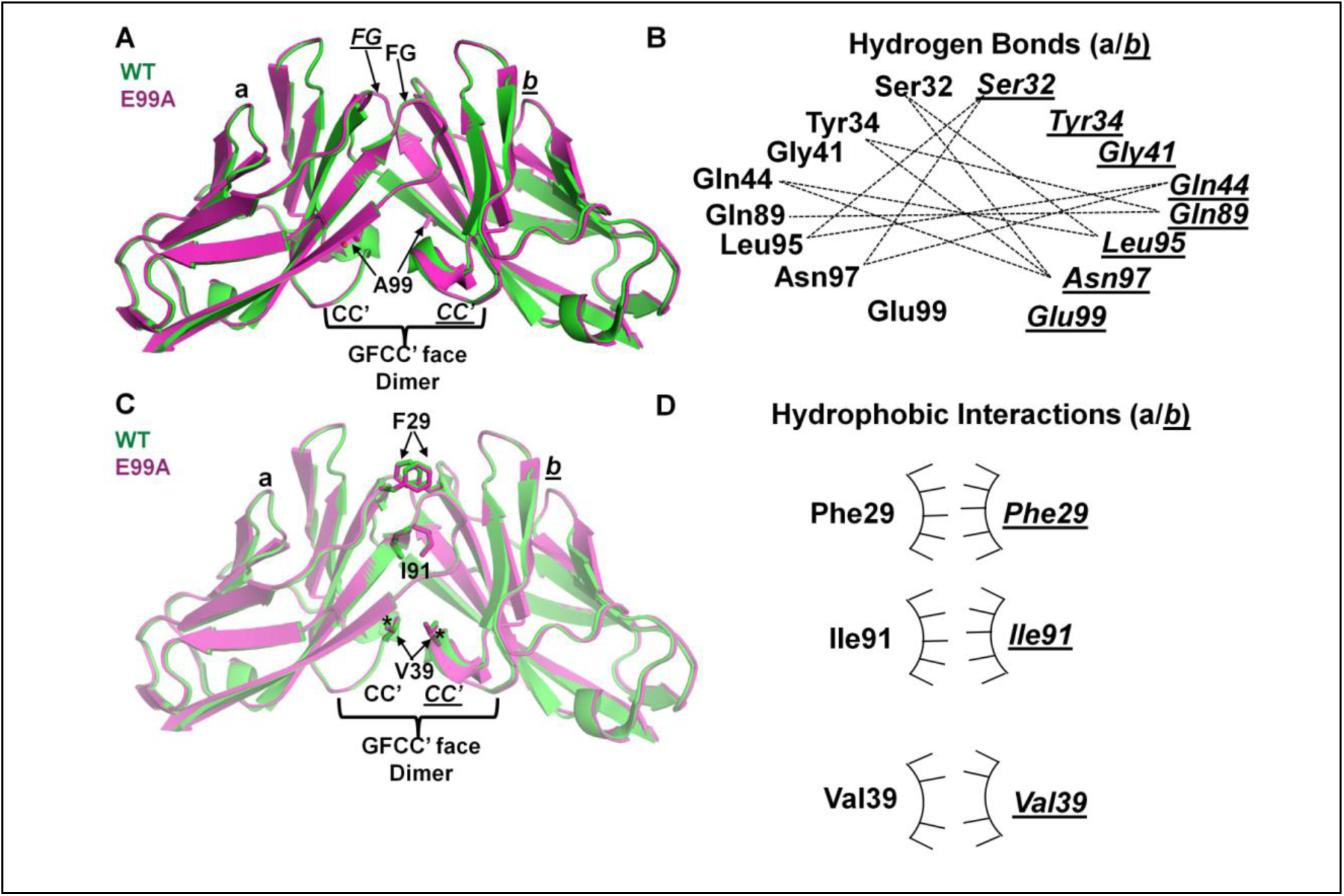
The E99A mutant crystal structure compared with the hCEACAM1 WT IgV (PDB code 4QXW) crystal structure. The labels, hydrogen bonds, and hydrophobic interactions using similar format as described in *Figure supplement 3* unless otherwise noted. (A) Ribbon diagram of the E99A mutant (magenta) and WT (green) IgV molecules (a) and (*b*) superimposed on each other. The stick representation of A99 residues as observed in the E99A crystal structure. (B) The hydrogen bonded interactions (dashed lines) between the residues described in Figure 2D, which mediate formation of the GFCC’ face E99A dimer. (C) The superimposition of the E91A mutant (magenta) and WT (green) IgV molecules (a) and (*b*), where residues F29, I91 and V39 make hydrophobic interactions. The V39 residues in E99A crystal structure make weaker hydrophobic interactions as evidenced by higher distance (3.9 Å) between V39-β carbons (marked by asterisk) compared to 3.7 Å in the WT. (D) The arc/stick representation of hydrophobic interactions by F29, I91 and V39 residues as observed in the E19A crystal structure. The weaker hydrophobic interactions mediated between two V39 residues are shown by fewer pointers on the hydrophobic arc.

**Figure supplement 4.**
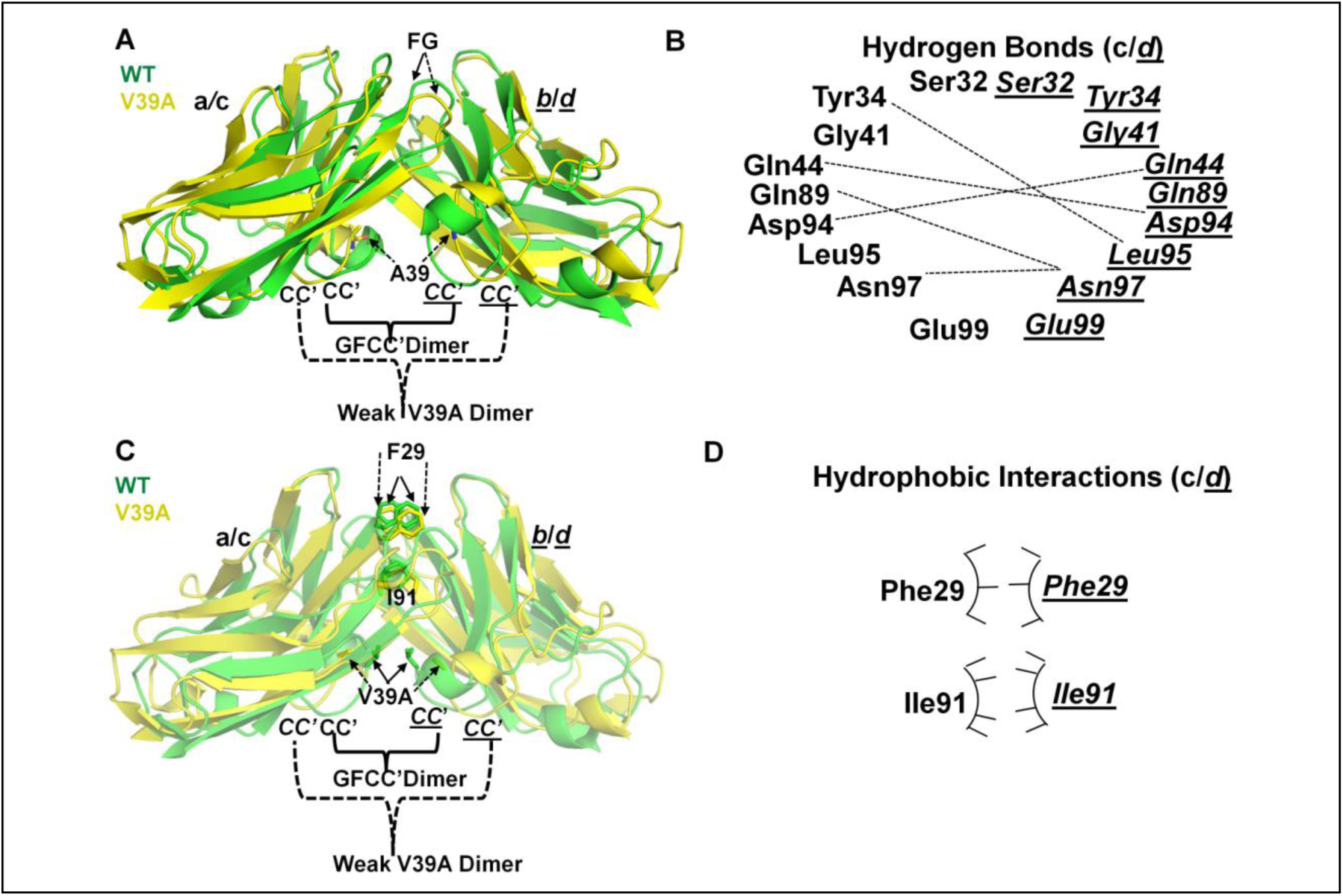
The structural basis of weak V39A mutant GFCC’ face dimer formation and comparison with hCEACAM1 WT IgV (PDB code 4QXW) crystal structure. (A) The structural superimposition of WT IgV molecules (a, b, colored green) and V39A mutant molecules (c, *d*, colored yellow), whereas CC’ and FG loops (dashed arrows) in yellow shows large conformational differences in the V39A crystal structure compared to CC’ and FG loops (solid arrows) of hCEACAM1 WT IgV (PDB code 4QXW) in green. The CC’ loops of WT dimer (PDB code 4QXW) are shown by solid arrows and mediates formation of GFCC’ face dimer The position of A39 residue on the CC’ loops is shown by stick representation, whereas this mutation results in weaker V39A dimer at the GFCC’ face due to the less numbered hydrogen bonded and weaker hydrophobic interactions, which is evident by increased distance between β carbon of A39 (10 Å) compared to distance of 3.7 Å between β carbon of V39 in the WT. (B) The fewer hydrogen bonded interactions that result in the weak V39A dimer formation between molecules (c) and (*d*) residues of the V39A mutant crystal structure are shown by dashed lines. The residues of molecules (c) and (*d*) are shown in bold and italics underlined, respectively. (C) The superimposition of the V39A mutant molecules (c, *d,* colored yellow) and WT molecules (a, *b*, colored green), whereas residues F29 and I91 in the yellow stick representation make weaker hydrophobic interactions in the weak V39A dimer. The A39 and V39 residues of the weak V39A dimer and WT dimer are shown by yellow stick/dashed arrows and green stick/solid arrows, representation, respectively. The CC’ loops are labeled as described above for WT and weak V39A dimer, and weaker hydrophobic interactions and conformation changes in the F29 residues of the weak V39A dimer are shown by dashed arrows compared to solid arrows in the WT. (D) The arc/stick representation of weaker hydrophobic interactions by F29, and I91 residues as observed between molecules (c) and (*d*) in the formation of weak V39 dimer of the V39A crystal structure. The weaker hydrophobic interactions mediated are shown by fewer pointers on the hydrophobic arc.

**Figure supplement 5.**
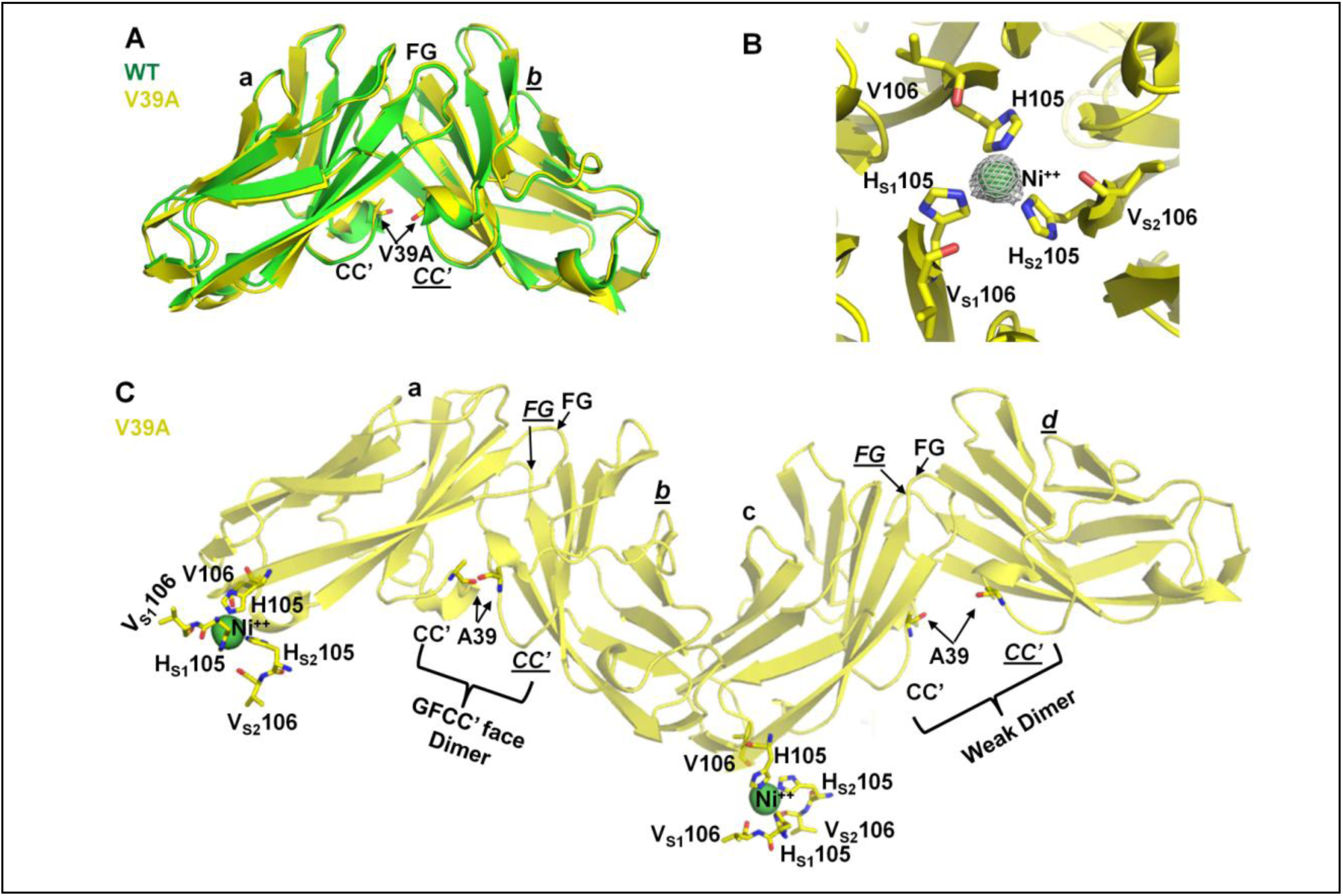
V39A mutant dimer formed by molecules (a) and (*b*) mimic hCEACAM1 WT GFCC’ face dimer (PDB code 4QXW) and residues H105 and V106 with their symmetry mates mediates binding with Nickel (Ni **^++)^**. (A) Ribbon diagram of the V39A mutant (yellow) and WT (green) IgV molecules (a) and (*b*) superimposed on each other and GFCC’C’ face dimer is shown. The CC’ and FG loops are labeled. (B) Interactions of six V39A mutant residues H105 and V106 from molecule (a) and its two symmetry mates 0100-100 (S1) and 02000000 (S2) with bound Ni **^++^** (green sphere) as observed in the V39A crystal structure. Residues H105, V106 from molecule (a), H_S1_105, V _S1_106, from symmetry molecule a_0100-100, and H_S2_105, V_S2_106 from symmetry molecule a_02000000, are highlighted in stick representation. These residues make interactions with Ni **^++^**, whereas nitrogen of three histidine rings of residues H105 (2.35 Å) and three carbonyl oxygen of residues V106 (4.53 Å) participates in hexa-coordinated interactions with Ni^++^. Similar interactions were also observed with molecule (c) and its symmetry mares in binding with second Ni^++^ as observed in the crystal structure. (C). Overall ribbon diagram of the four molecules (a, *b*, c, *d*) in yellow, whereas A39 residue of each molecule is shown by stick representation and binding of Ni^++^ with molecule (a) or (C) and their symmetry mates residues H105 and V106 as described above. Molecules (a) and (*b*) makes GFCC’C’ face dimer that mimics WT dimer. In contrast, molecules (c) and (*d*) make a weak V39A dimer where CC’ loops are apart and show large conformation differences compared to WT GFCC’ face dimer.

**Figure supplement 6.**
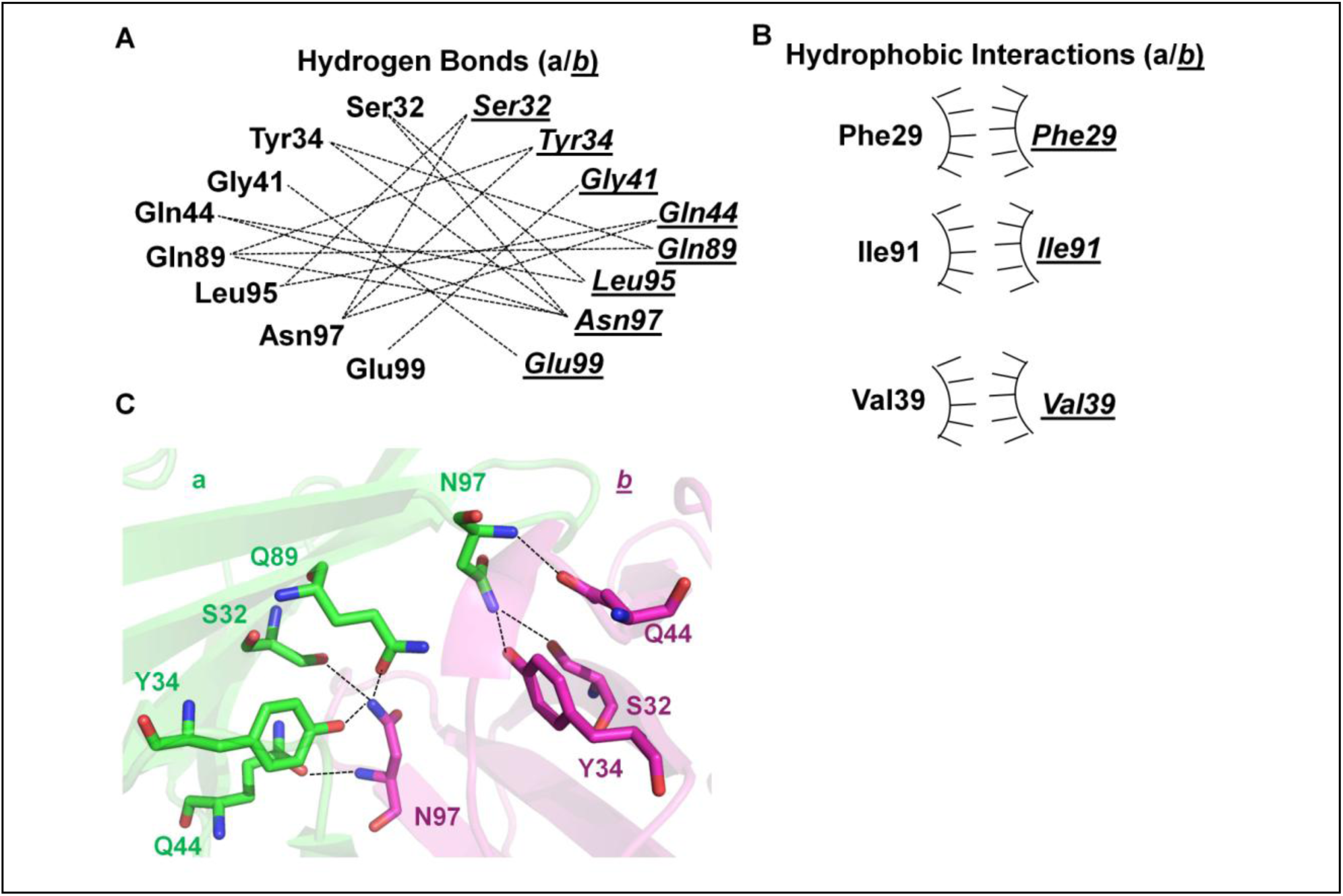
Hydrogen bonds and hydrophobic interactions of the IgV of hCEACAM1 homodimer (PDB code 4QXW) and N97-mediated asymmetrical hydrogen bonded interactions. (A) Hydrogen bonded interactions of GFCC’ face residues. The molecule (a) residues are labeled in bold, and molecule (*b*) residues are labeled in italics and underlined. The hydrogen bonded interactions across the GFCC’ face by residues described above are shown by dashed lines. (B) The hydrophobic interactions of GFCC’ face F29, I91, and V39 residues are shown by arc/point representations. The molecule (a) and (*b*) residues are labeled as described above. (C) Seven asymmetrical hydrogen bonded interactions (dashed lines) mediated by N97A residues of molecule a in red and molecule (*b*) in magenta as observed in the hCEACAM1 WT homodimer structure (PDB code 4QXW). The N97 residue from molecule (a) makes three hydrogen bonded interactions with residues S32, Y34, and Q44 of molecule (*b*) and N97 residue from molecule (*b*) makes four hydrogen bonded interactions with residues S32, Y34, Q44, and Q89 of molecule (a).

**Figure supplement 7.**
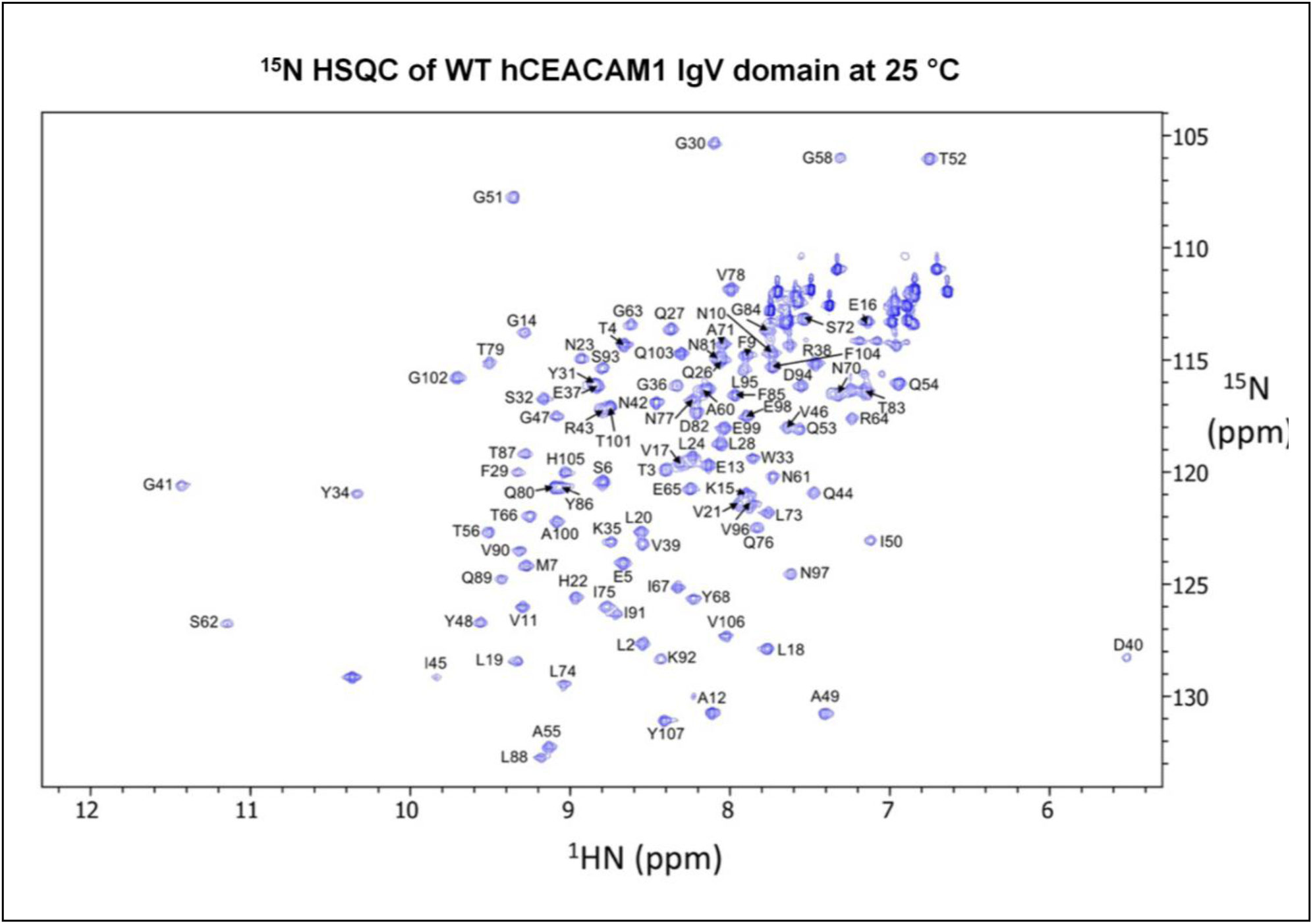
^15^N HSQC spectra of hCEACAM1 IgV WT and N97A mutant proteins. (A) Assigned ^15^N-HSQC spectrum of hCEACAM1 IgV WT protein (blue).

**Figure supplement 7B.**
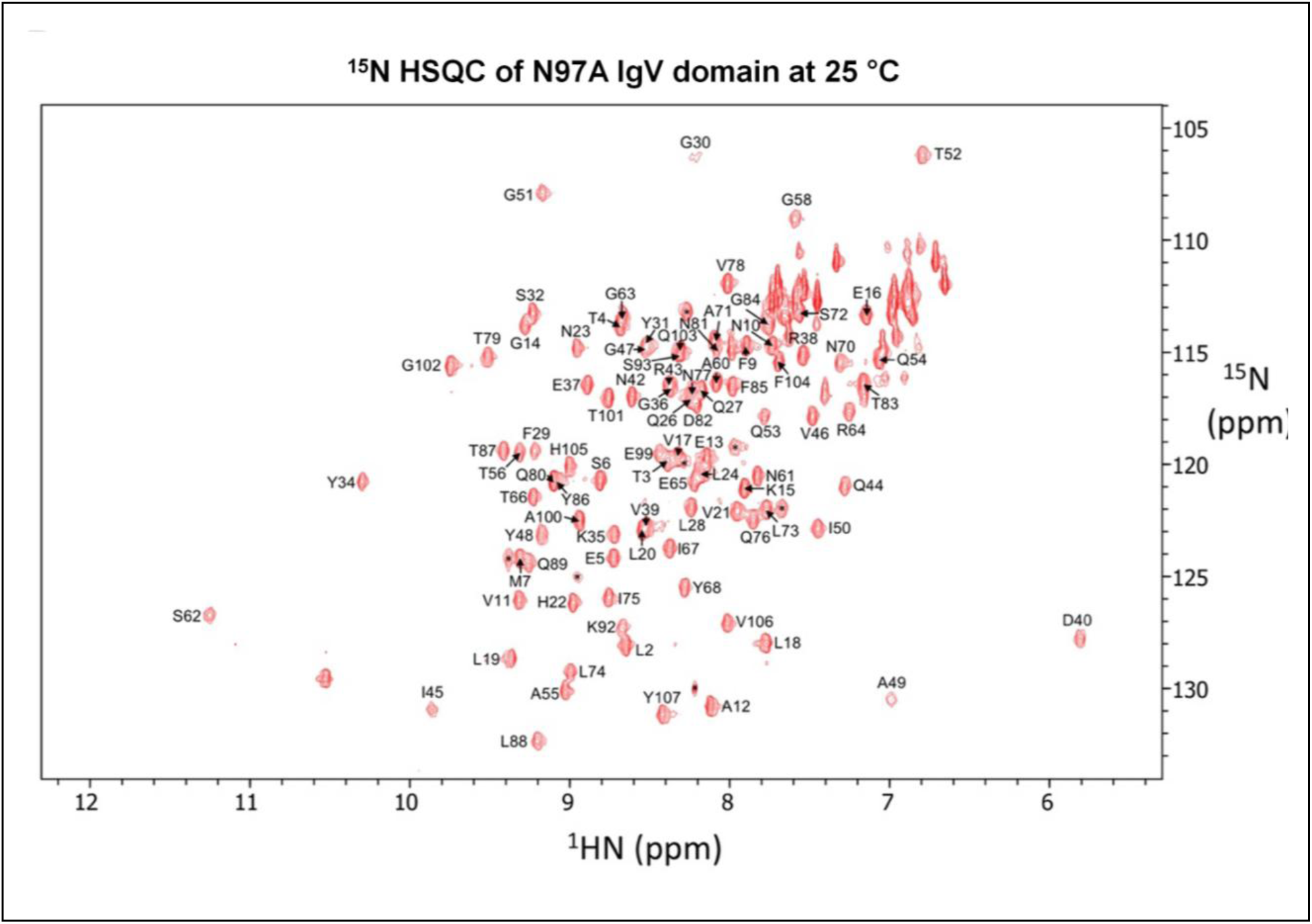
Assigned ^15^N-HSQC spectrum of N97A mutant hCEACAM1 (red) IgV protein.

**Figure supplement 8.**
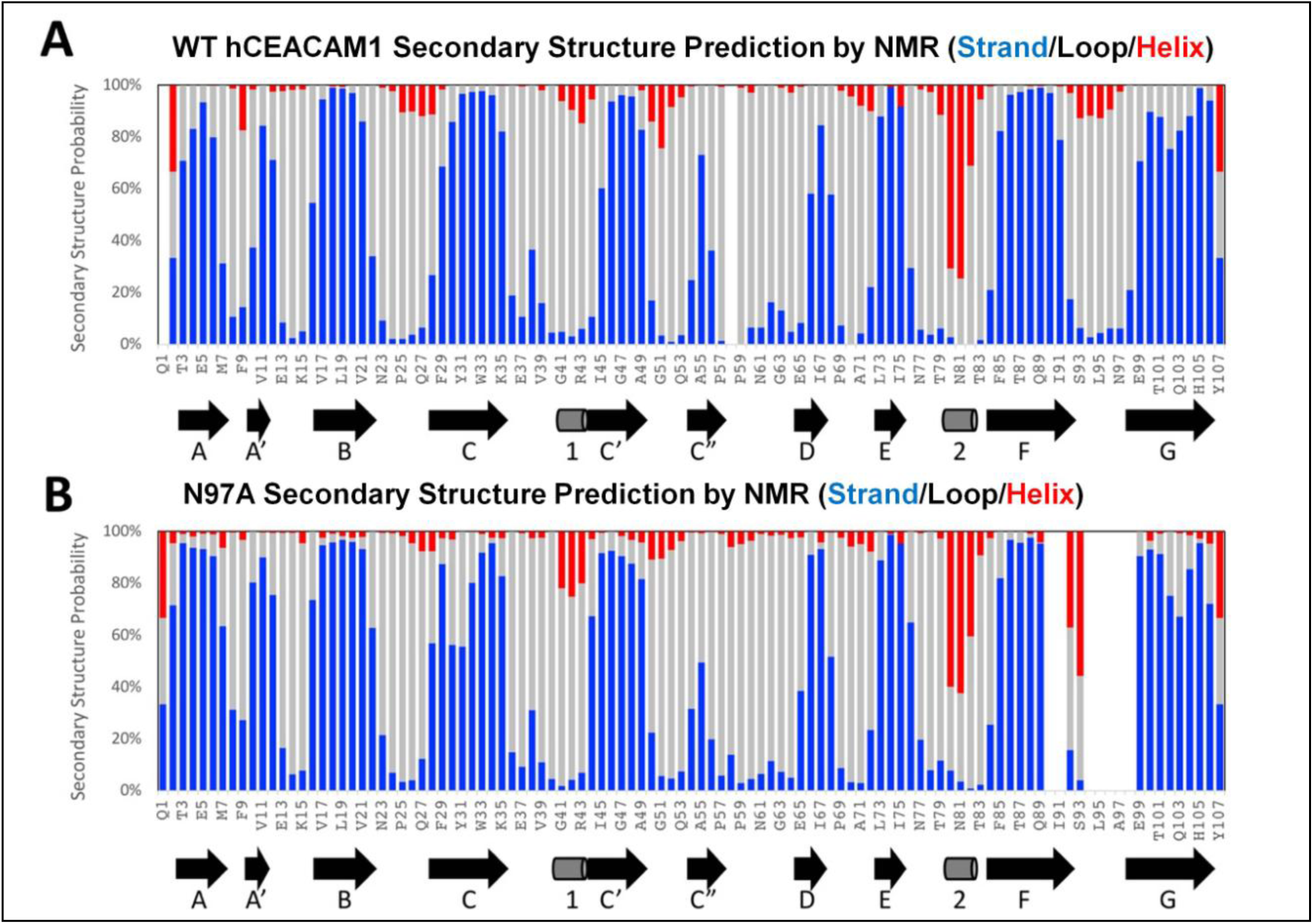
Secondary structures prediction and comparison. (A) Secondary structures of hCEACAM1 WT IgV predicted from NMR chemical shift values, in comparison with the secondary structure elements from the X-ray structure (PDB code 4QXW) depicted below. (B) Secondary structures of N97A mutant hCEACAM1 IgV predicted from NMR chemical shift values, in comparison with the secondary structure elements from the X-ray structure (PDB code 6XO1) depicted below. The probabilities of predicted beta-strand, loop, and alpha-helix are colored in blue, grey, and red respectively.

**Figure supplement 9.**
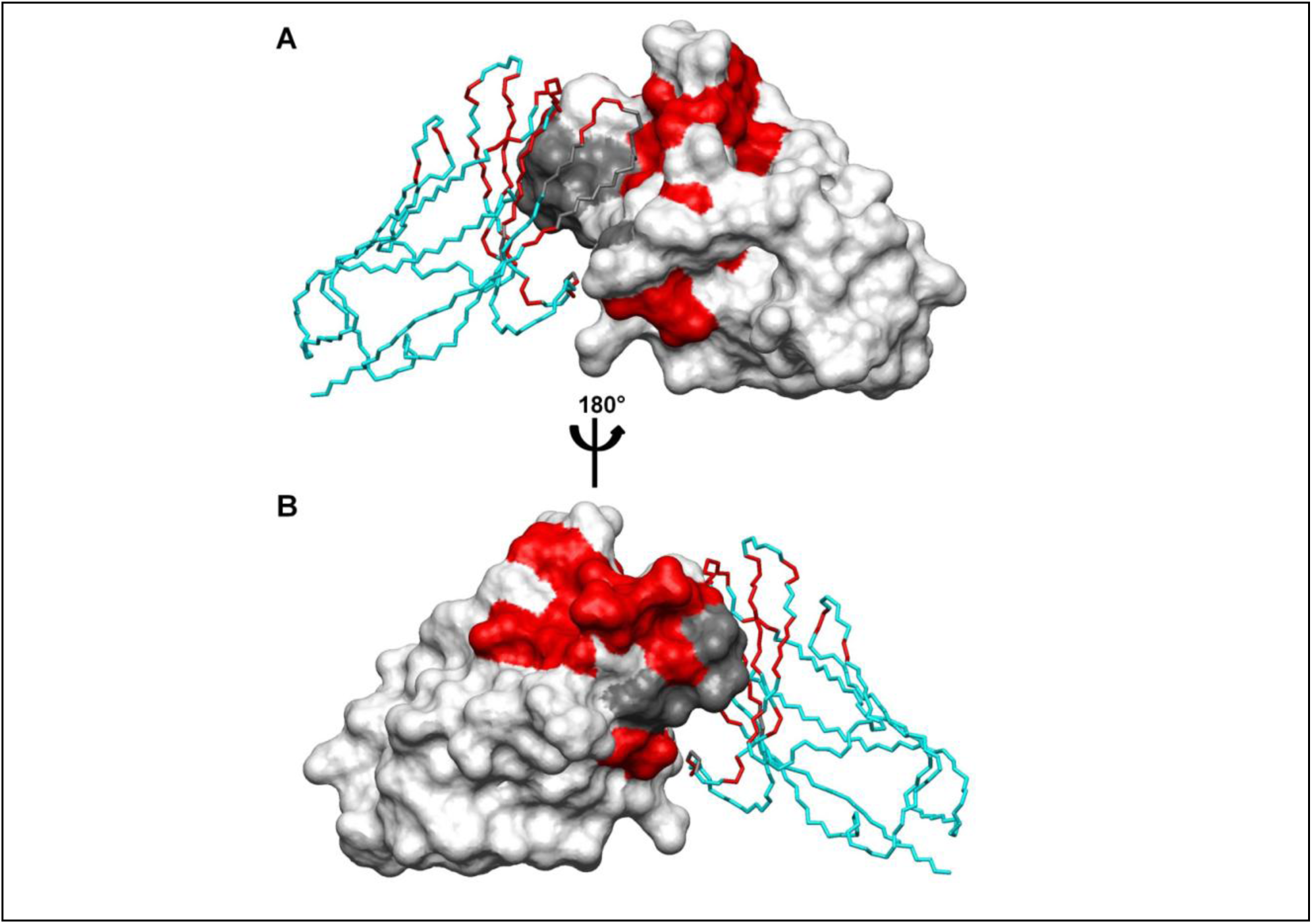
Mapping of NMR chemical shift changes. (A) Front and (B) back view of the crystal structure of the hCEACAM1 WT IgV dimer (PDB code 4QXW) with one unit shown in molecular surface representation (white) and another in alpha carbon traces (cyan). The residues with combined NMR chemical shift changes larger than 0.2ppm are colored in red, while residues unassigned in N97A mutant hCEACAM1 IgV are colored in grey.

**Figure supplement 10.**
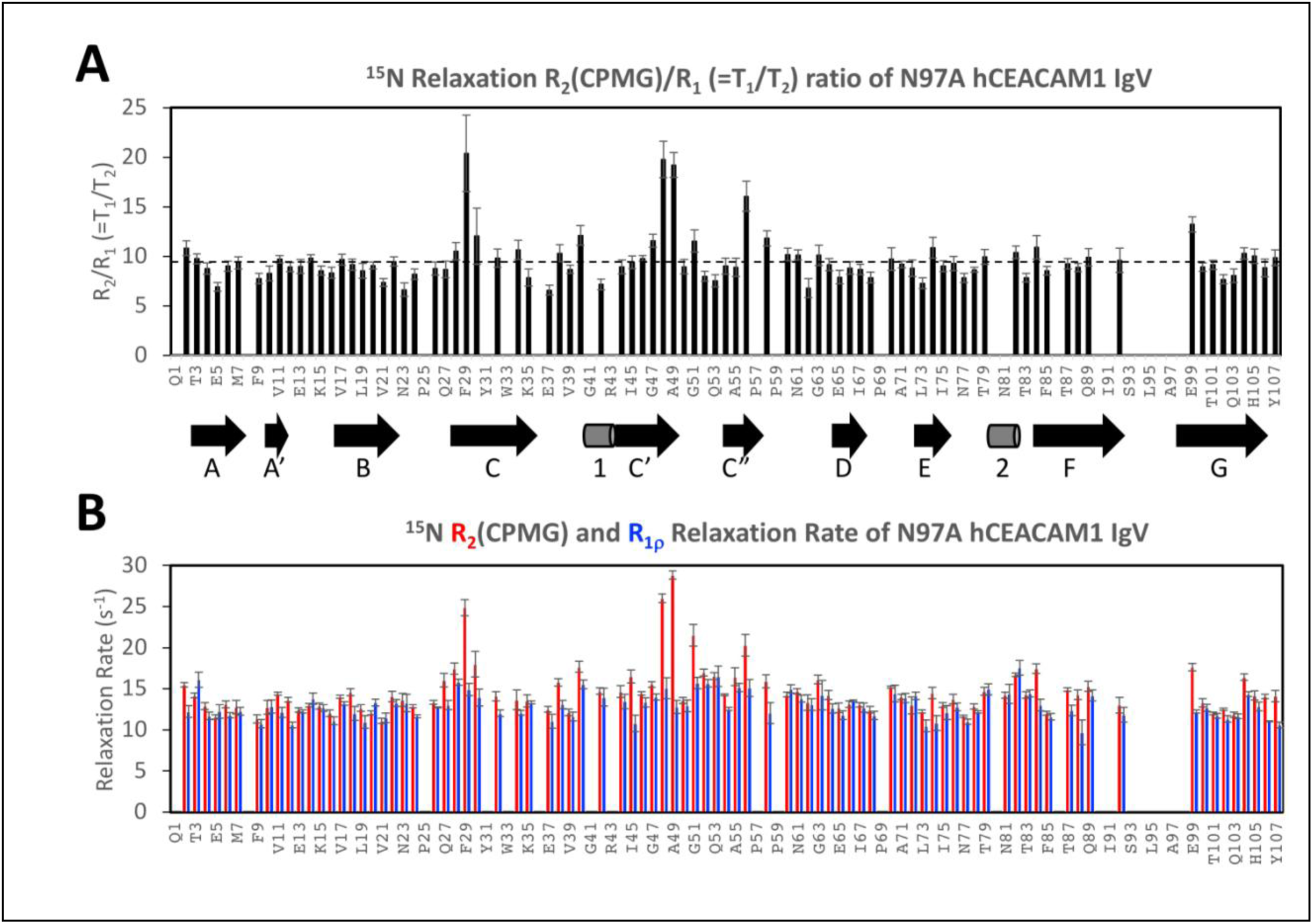
NMR relaxation rates. (A) The ratio of NMR relaxation rates R_2_/R_1_ (same as the ratio of relaxation times T_1_/T_2_) of N97A mutant hCEACAM1 IgV showing remarkable dynamics of several residues at the GFCC’ face. T_2_ was measured by a CPMG type experiment (with τ = 625μs). (B) The comparison between R_2_ (red) and R_1rho_ (blue) NMR transverse relaxation rates of N97A mutant hCEACAM1 IgV showing chemical exchange effects that are better suppressed in the T_1rho_ experiment (with a field-locking strength of 1.5kHz) than in the T_2_ CPMG experiment (with a CPMG τ = 625μs).

**Figure supplement 11.**
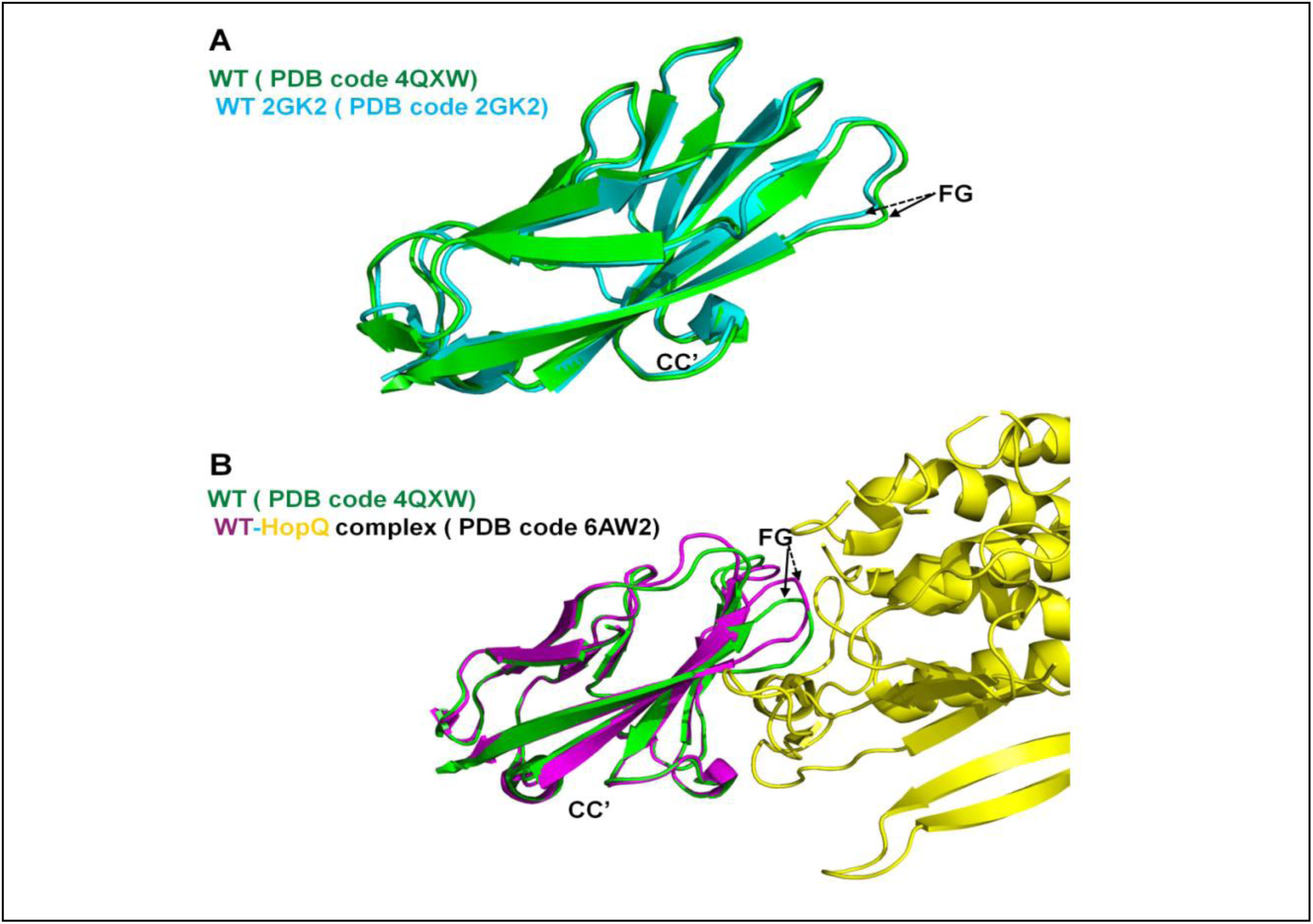
The conformational flexibility of the hCEACAM1 WT GFCC’ face. (A) Structural alignment of a hCEACAM1 molecule of WT homodimer in green (PDB code 4QXW) with a hCEACAM1 molecule of WT crystal structure in cyan (PDB code 2GK2) by ribbon representation. CC’ and FG loops are labeled, where conformational differences of FG loops in two crystal structures are indicated by solid and dashed lines. (B) Structural alignment of a hCEACAM1 molecule of WT homodimer in green (PDB code 4QXW) with a hCEACAM1 molecule of WT-HopQ complex crystal structure in magenta (PDB code 6AW2). For clarity, only half of the HopQ molecule (yellow) is shown. CC’ and FG loops are labeled as described above.

**Figure supplement 12.**
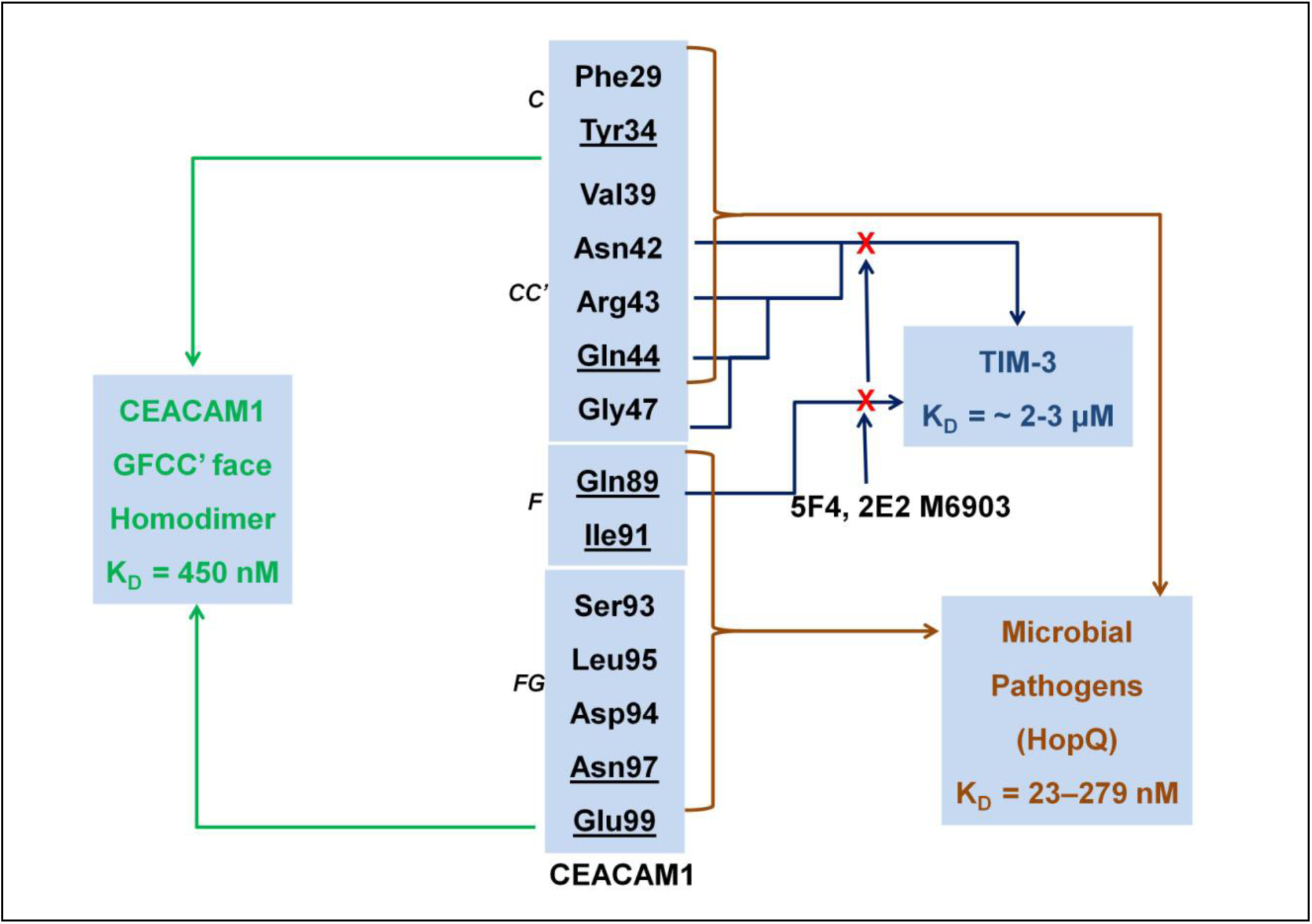
The GFCCC” face residues network of hCEACAM1 mediates homodimer formation and interactions with various ligands including hTIM-3 and microbial pathogens such as HopQ. The GFCCC” face residues F29, Y34,G41,Q44, Q89, I91, L95,N97 and E99 make hydrophobic and hydrogen bonded interactions with 450 nM affinity (K_D_) of homodimer formation. The residues of GFCCC” face also participate in the interactions with hTIM-3 to regulate immune tolerance and exhaustion in which residues N42, R43, Q44, G47 and Q89 mediate the binding with hCEACAM1 as shown by site directed mutagenesis studies. The binding between hCEACAM1 and hTIM-3 as shown by NMR, SPR and ELISA exhibits a K_D_ of ∼2-3 μM with blockade of this interaction by 5F4 (anti-hCEACAM1 monoclonal), 2E2 (anti-hTIM3 monoclonal antibody) and M6903 (anti-hTIM3 monoclonal antibody), and anti-TIM3 polyclonal antibodies. Further, microbial pathogens such as HopQ, OPA proteins and Afa/Dr adhesins also target this GFCC’ face for cellular invasion and immune evasion. Specially, the HopQ GFCC’ face residues F29, V39, Q44, Q89, I91, V96 binds hCEACAM1 with higher K_D_ of 23-279 nM than K_D_ of homodimer formation and abrogate GFCC’ face.

**Figure supplement 13.**
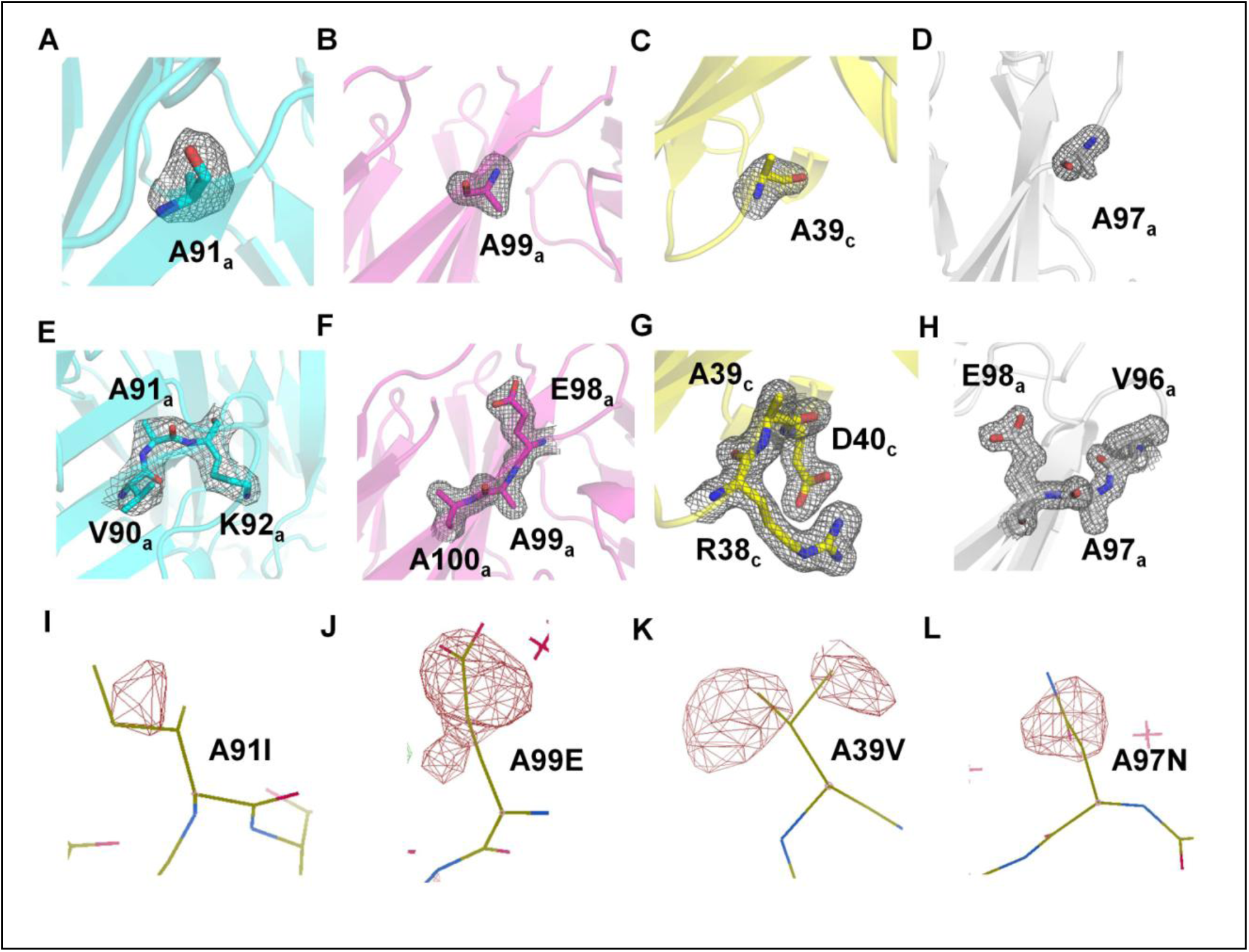
Electron density maps of hCEACAM1 IgV I91A, E99A, V39 and N97A mutants. (A-D) Fo-Fc map contoured at 3.0 σ level of hCEACAM1 mutants. Map is derived from initial model where residues A91 (panel A), A99 (panel B), A39 (panel C), and A97 (panel D) were not fitted and significant positive Fo-Fc electron density at 3.0 σ level was observed. Figures A-D showing superimposition of the Fo-Fc map on the final refined model and residues A91 from molecule a (panel A), A99 from molecule a (panel B), A39 from molecule c (panel C), and A97 from molecule a (panel D) are shown by stick representations. The molecule name for the identification of these residues is shown in the subscript. (E-H) 2Fo-Fc map at 1.0 σ level of hCEACAM1 mutants. Map is derived from final refined model where residues V90-A91-K92 from molecule a (panel E), E98-A99-A100 from molecule a (panel F), R38-A39-D40 from molecule c (panel G), and V96-A97-E98 from molecule a (panel H) are fitted to the initial model after many rounds of model building and refinement. Figures E-H showing superimposition of 2Fo-Fc map on the final refined model and residues are shown by stick representations. (I-L) Further validation of hCEACAM1 IgV I91A, E99A, V39 and N97A mutants. The final refined model of each mutant was reverse mutated to the respective residue present in the WT, whereas A91I (panel I), A99E (panel J), A39V (panel K), and A97N (panel L) mutations were done in the I91A, E99A, V39 and N97A mutants coordinates, respectively and one round of refinement cycle was performed. The observed negative Fo-Fc map at 3.0 σ level for each mutant clearly shows negative density around reverse mutation site and validates proper model building and refinement of the hCEACAM1 mutants.

## Supplement Tables

**Supplementary Table 1:**
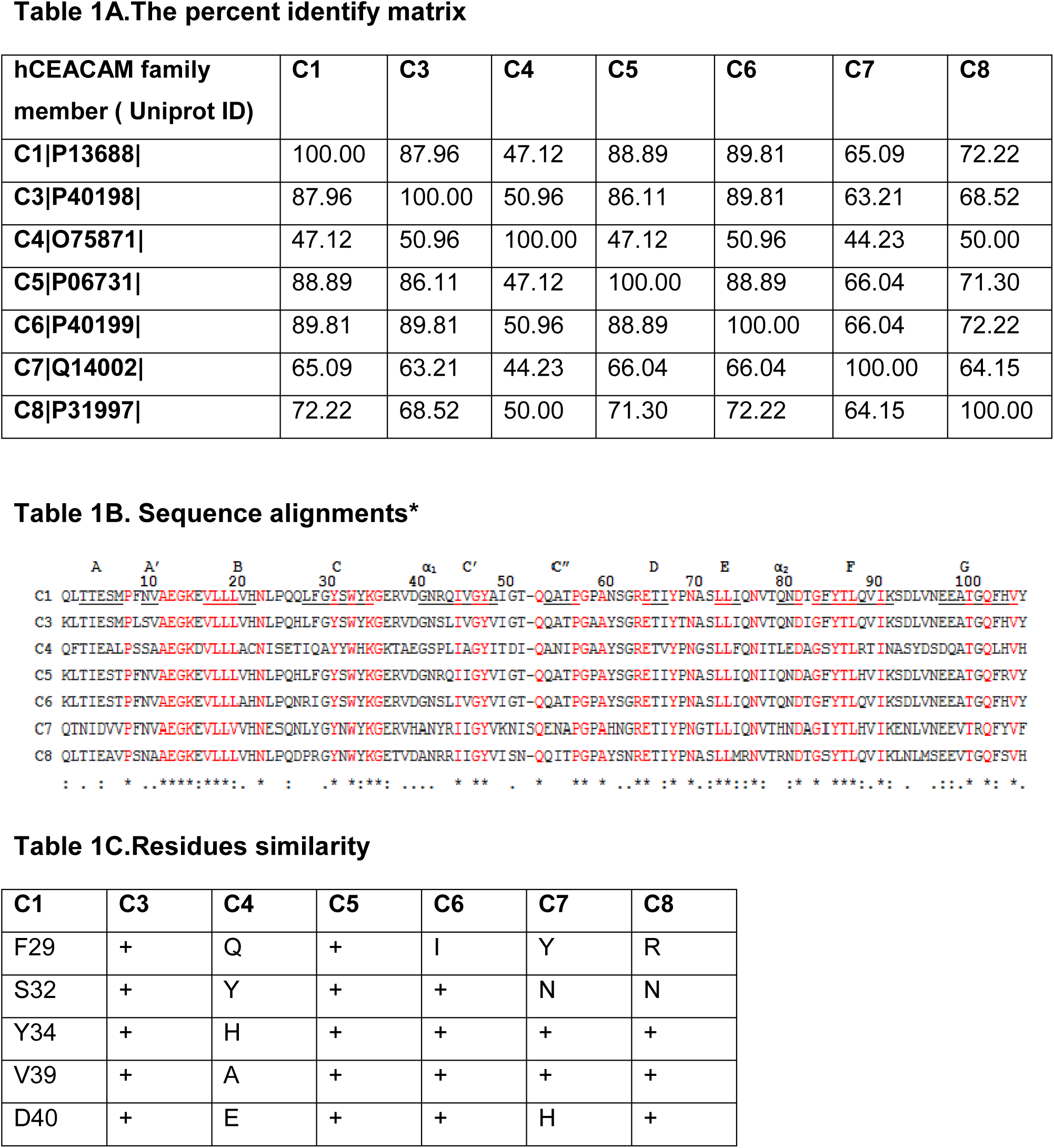
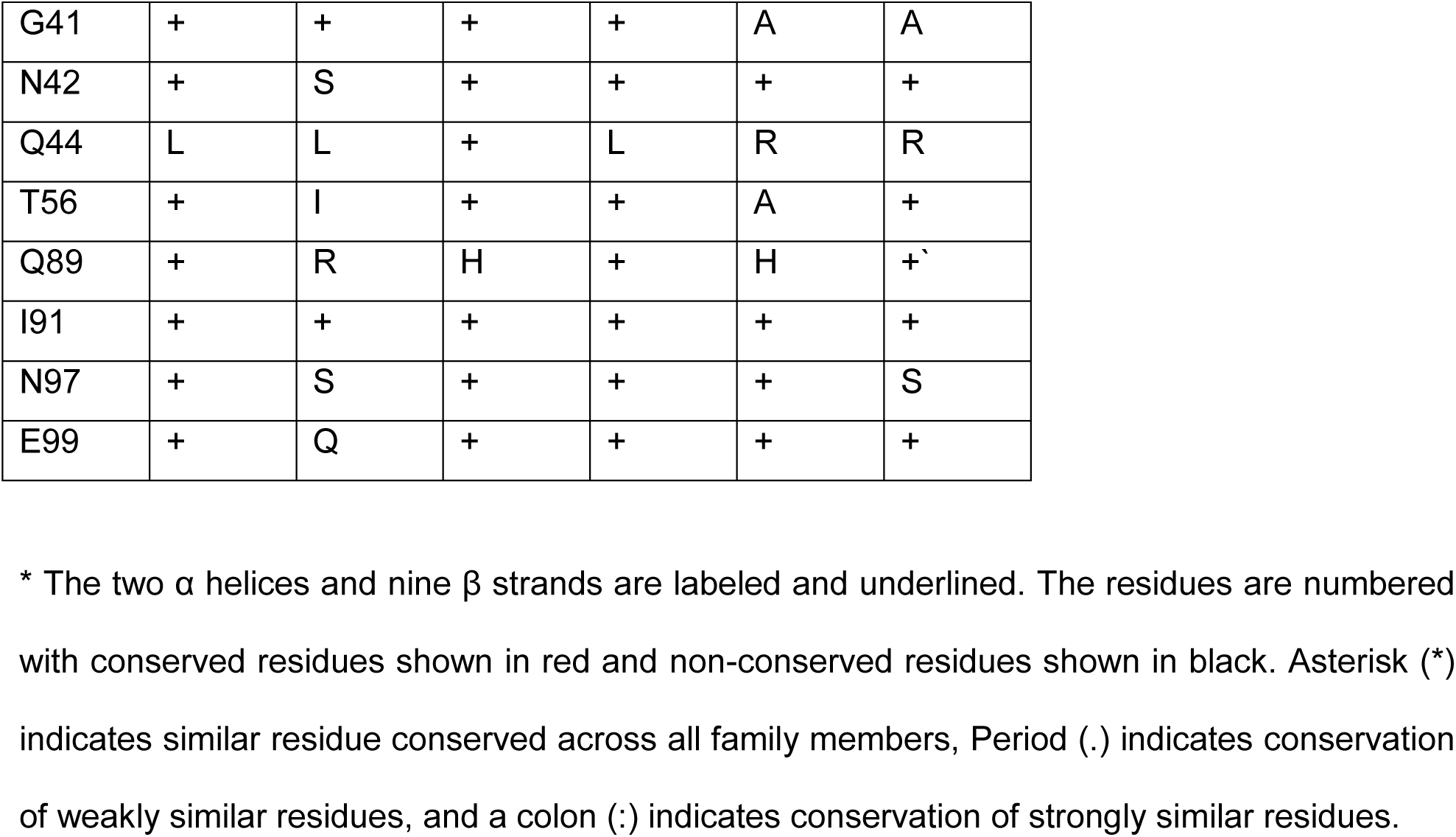
The percent identify matrix, sequence alignments and residues similarity of IgV of the Human CEACAM family members including CEACAM1 (C1), CEACAM3 (C3), CEACAM4 (C4), CEACAM5 (C5), CEACAM6 (C6), CEACAM7 (C7) and CEACAM8 (C8).

**Supplementary Table 2:** Hydrogen bonded interactions as observed in human CEACAM1 WT dimer (PDB code 4QXW) and V39A (PDB code 6XNW), I91A (PDB code 6XNT), N97A (PDB code 6XO1), E99A (PDB code 6XNO) crystal structures. Table 2A: Human CEACAM1 WT dimer, molecules a/*b*

| Interactions | Molecule (a) residues<br>( Interacting atom) | Molecule (b) residues<br>( Interacting atom) | Distance (Å) |
| --- | --- | --- | --- |
| 1 | L95[ O ] | S32[ OG ] | 2.96 |
| 2 | E99[ OE2] | G41[ N ] | 2.88 |
| 3 | L95[ O ] | Q44[ NE2] | 2.80 |
| 4 | Y34[ OH ] | Q89[ NE2] | 3.35 |
| 5 | Q89[ OE1] | Q89[ NE2] | 3.06 |
| 6 | Q44[ OE1] | N97[ N ] | 2.96 |
| 7 | S32[ OG ] | N97[ ND2] | 2.94 |
| 8 | Y34[ OH ] | N97[ ND2] | 3.03 |
| 9 | Q89[ OE1] | N97[ ND2] | 3.82 |
| 10 | S32[ OG ] | L95[ O ] | 2.90 |
| 11 | G41[ N ] | E99[ OE1] | 2.69 |
| 12 | Q44[ NE2] | L95[ O ] | 2.80 |
| 13 | Q89[ NE2] | G89[ OE1] | 2.98 |
| 14 | Q89[ NE2] | Y34[ OH ] | 3.58 |
| 15 | N97[ N ] | Q44[ OE1] | 3.21 |
| 16 | N97[ ND2] | S32[ OG ] | 3.06 |
| 17 | N97[ ND2] | Y34[ OH ] | 3.46 |

| Interactions | Molecule (a) residues<br>( Interacting atom) | Molecule (b) residues<br>( Interacting atom) | Distance (Å) |
| --- | --- | --- | --- |
| 1 | L95[ O ] | S32[ OG ] | 2.94 |
| 2 | E37[ O ] | R38[ NH1] | 3.11 |
| 3 | E99[ OE1] | G41[ N ] | 2.67 |
| 4 | L95[ O ] | Q44[ NE2] | 3.07 |
| 5 | Q89[ OE1] | Q89[ NE2] | 3.05 |
| 6 | Q44[ OE1] | N97[ N ] | 3.05 |
| 7 | S32[ OG ] | N97[ ND2] | 3.35 |
| 8 | S32[ OG ] | L95[ O ] | 2.95 |
| 9 | R38[ NH1] | E37[ O ] | 3.64 |
| 10 | G41[ N ] | E99[ OE1] | 2.70 |
| 11 | Q44[ NE2] | L95[ O ] | 2.66 |
| 12 | Q89[ NE2] | Y34[ OH ] | 3.39 |
| 13 | Q89[ NE2] | Q89[ OE1] | 3.17 |
| 14 | N97[ N ] | Q44[ OE1] | 3.05 |
| 15 | N97[ ND2] | S32[ OG ] | 3.29 |
| 16 | N97[ ND2] | Y34[ OH ] | 3.34 |

| Interactions | Molecule (c) residues<br>( Interacting atom) | Molecule (d) residues<br>( Interacting atom) | Distance (Å) |
| --- | --- | --- | --- |
| 1 | Q89[ OE1] | N 97[ ND2] | 2.91 |
| 2 | D 94[ O ] | Q44[ NE2] | 3.19 |
| 3 | N97[ OD1] | N97[ ND2] | 2.45 |
| 4 | Y34[ OH ] | L95[ O ] | 3.85 |
| 5 | Q44[ NE2] | D94[ O ] | 2.97 |

| Interactions | Molecule (a) residues<br>( Interacting atom) | Molecule (b) residues<br>( Interacting atom) | Distance (Å) |
| --- | --- | --- | --- |
| 1 | L95[ O ] | S32[ OG ] | 3.11 |
| 2 | E99[ OE1] | G41[ N ] | 3.17 |
| 3 | L95[ O ] | Q44[ NE2] | 2.76 |
| 4 | Q89[ OQE1] | Q89[ NE2] | 2.93 |
| 5 | Q89[ OE1] | N97[ ND2] | 3.44 |
| 6 | S32[ OG ] | L95[ O ] | 2.57 |
| 7 | G41[ N ] | E99[ OE1] | 2.88 |
| 8 | Q44[ NE2] | L95[ O ] | 3.04 |
| 9 | Q89[ NE2] | Q89[ OE1] | 2.84 |
| 10 | N 97[ N ] | Q44[ OE1] | 3.81 |
| 11 | N97[ ND2] | S32[ OG ] | 3.86 |
| 12 | N97[ ND2] | Q89[ OE1] | 3.63 |

| Interactions | Molecule (a) residues<br>( Interacting atom) | Molecule (b) residues<br>( Interacting atom) | Distance (Å) |
| --- | --- | --- | --- |
| 1 | N26[ OE1] | I67[ N ] | 2.88 |
| 2 | N26[ NE2] | I67[ O ] | 2.94 |
| 3 | N53[ NE2] | G58[ O ] | 3.89 |
| 4 | N53[ NE2] | N61[ O ] | 3.51 |

| Interactions | Molecule (a) residues<br>( Interacting atom) | Molecule (b) residues<br>( Interacting atom) | Distance (Å) |
| --- | --- | --- | --- |
| 1 | S32[ OG ] | N97[ ND2] | 3.04 |
| 2 | Y34[ OH ] | N97[ ND2] | 3.35 |
| 3 | Y34[ OH ] | Q89[ NE2] | 3.57 |
| 4 | Q44[ OE1] | N97[ N ] | 3.11 |
| 5 | Q89[ OE1] | Q89[ NE2] | 3.25 |
| 6 | L95[ O ] | S32[ OG ] | 2.89 |
| 7 | L95[ O ] | Q44[ NE2] | 2.84 |
| 8 | S32[ OG ] | L95[ O ] | 3.00 |
| 9 | Q 44[ NE2] | L95[ O ] | 2.78 |
| 10 | Q89[ NE2] | Q89[ OE1] | 3.06 |
| 11 | N97[ N ] | Q44[ OE1] | 3.42 |
| 12 | N97[ ND2] | S32[ OG ] | 3.35 |

**Supplementary Table 3:** PDB PISA complex significance score (CSS) analysis of human CEACAM1 WT dimer (PDB code 4QXW) and V39A (PDB code 6XNW), I91A (PDB code 6XNT), N97A (PDB code 6XO1), E99A (PDB code 6XNO) crystal structures.

| Crystal structure | CSS Score |
| --- | --- |
| CEACAM1 WT ( molecules *a/b*) | 0.9 |
| V39A ( molecules *a/b*) | 1.0 |
| V39A ( molecules *c/d*) | 0.0 |
| I91( molecules *a/b*) | 1.0 |
| N97A ( molecules *a/b*) | 0.0 |
| E99A ( molecules *a/b*) | 0.63 |

